# Cell-to-cell variability in the yeast pheromone response: high throughput screen identifies genes with different effects on transmitted signal and response

**DOI:** 10.1101/093187

**Authors:** C. G. Pesce, W. Peria, S. Zdraljevic, D. Rockwell, R. C. Yu, A. Colman-Lerner, R. Brent

**Affiliations:** Fred Hutchinson Cancer Research Center, Seattle, Washington 98109; IFIBYNE-UBA-CONICET and D, Facultad de Ciencias Exactas y Naturales, Universidad de Buenos Aires, Buenos Aires C1428EHA, Argentina; The former Molecular Sciences institute, Berkeley, California 94704

## Abstract

Populations of isogenic cells often respond coherently to signals despite differences in protein abundance and cell state. Our previous work in the *Saccharomyces cerevisiae* pheromone response system (PRS) uncovered processes that reduced cell-to-cell variation in signal and response. To understand these and other processes that controlled variation, we generated a whole-genome collection of haploid strains with deletions in non-essential genes and used high-throughput flow cytometry to screen more than 1000. We identified 50 “variation genes” required for normal cell-to-cell variability in signal and response. Some genes affected only signal variability, signal strength, or system output, defining these quantities as separable “axes” of system behavior. Two genes affected cytoplasmic microtubule function.

## Introduction

Cell signaling systems transmit information about the external environment, enabling cells to respond to extracellular signals. Although much is known about the operation of these systems, however, the means they use to ensure precise and accurate signal transmission and cell response remain largely unknown. Moreover, in metazoan tissues, populations of cells must determine concentrations of extracellular signaling and make appropriate fate decisions in response to those determinations. Coherence in these cell population responses is critical for the choreographed sequence of cell and tissue interactions during embryonic development, and for regulated cell division and differentiation during tissue maintenance in the adult.

Yet cells, even genetically identical cells in common environments behave differently. Such variability was first demonstrated in studies of bacteria and phages. In 1945 Delbrück showed large cell-to-cell variation in the yield of T1 phage from individual *Escherichia coli* cells (Delbrück, 1945), and Lieb demonstrated a reproducible binary distribution of clonal *E. coli* either being lysed or lysogenized after infection with phage λ (Lieb, 1953). Later, Novick and Weiner showed that the time of induction of the *lac* operon in single cells was highly variable (Novick and Weiner, 1957), and Spudich and Koshland found persistent non-genetic cell-to-cell differences in chemotactic behavior of individual genetically identical *Salmonella typhimurium* bacteria (Spudich and Koshland, 1976). More recently, cell-to-cell variation in clonal behavior has been shown in mammalian cells; for example, during the differentiation of hematopoietic progenitor cells into erythroid and myeloid cells (Chang et al., 2008), the decision to die in response to a pro-apoptotic drug (Sigal et al.; Spencer et al., 2009) and the activation of latent HIV proviruses (Weinberger et al., 2005).

We and others have studied cell-to-cell variability in the cell fate decision system that controls mating in *S. cerevisiae*, the pheromone response system (PRS) (Colman-Lerner et al., 2005; Elf et al.; Novick and Weiner, 1957; Paliwal et al., 2007;Ricicova et al., 2013; Yu et al.). The PRS is prototypic, in that it uses a GPCR, whose G protein couples in turn to a scaffolded-MAPK cascade (Dohlman and Thorner, 2001) (Figure 1a). Sensing of pheromone concentration via the receptor causes outputs including induction of genes at appropriate levels (here called “system output”) that depend on a set of proteins here called the signaling arm of the PRS. Determination of the direction of a gradient of pheromone concentration, and subsequent growth towards a mating partner, depend on proteins here called the polarity determination arm of the system. Our previous work quantified system output by expression from PRS responsive and control reporter genes (see Box). It separated the cell-to-cell variation in output into two contributions. The first of these was from events upstream of the promoter, affecting a signal transmission or “pathway” subsystem, P, quantified as η^2^(P). The second contribution was from variation affecting either pre-existing cell-to-cell differences in capacity of a “gene expression” subsystem η^2^(G) or rapid-acting changes in gene expression “intrinsic noise”, or η^2^(γ). This work further established analytically that cell-to-cell differences in (P) were caused by η^2^(L), (differences in L, the capacity component of the signal transmission subsystem at the start of the experiment) and η^2^(λ), rapid acting changes in signal during the measurement, but we could not separate η^2^(L) and η^2^(λ) experimentally.

**Figure 1.**
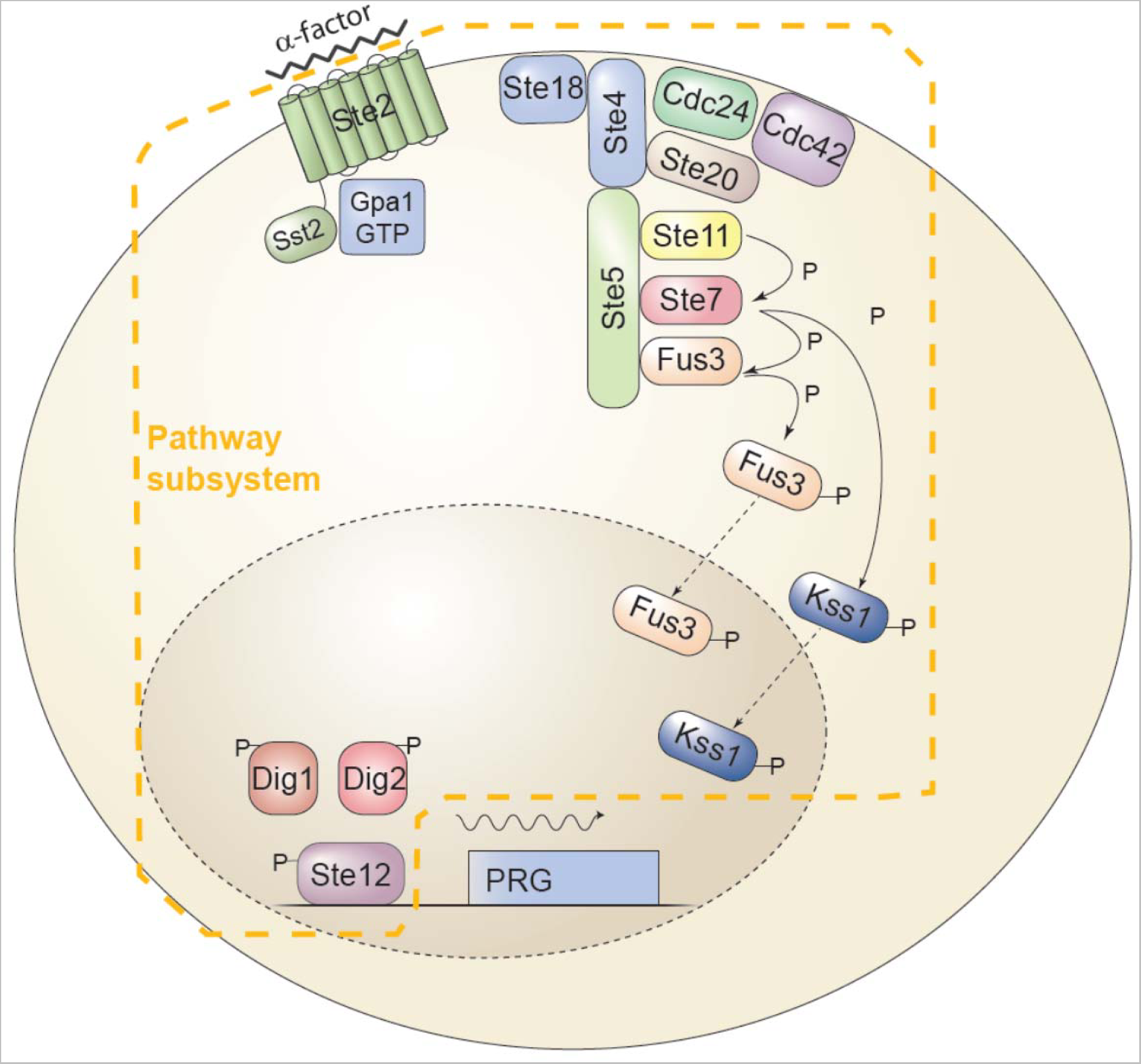
The yeast Pheromone Response System (PRS). **A.** The signaling arm of the yeast pheromone response system (PRS) and the analytical framework used to quantify variation in its operation. Haploid yeast cells react to pheromone secreted from cells of the opposite mating type with a series of cellular responses, including cell cycle arrest and induction of gene expression, that eventually lead the conjugating cells to form a diploid. Signaling in the PRS is system is prototypic of many eukaryotic signal transduction pathways. A seven-helix transmembrane receptor, Ste2 in the MATa cell depicted, causes the dissociation of the α subunit of a trimeric G-protein, Gpa1, from the βγ dimer, Ste4/Ste18. This event causes the recruitment to the plasma membrane of the scaffold protein Ste5, leading to the assembly and activation of the MAP kinase cascade (MAPKKK Ste11, MAPKK Ste7 and the Erk1/2-like MAPKs Fus3 and Kss1 (Dohlman and Thorner, 2001). In the cytoplasm, activated Fus3 and Kss1 regulate targets including Ste5 (Bush and Colman-Lerner, 2013; Yu et al., 2008), and in the nucleus, they activate Ste12 (Tedford et al., 1997). These events comprise the Pathway subsystem, P; i.e. the subsystem that transmits the signal to the promoters of inducible genes. Activation of Ste12 leads to the induction of approximately 100 pheromone responsive genes (PRGs) (Roberts et al., 2000) and their expression via the Expression subsystem E (defined in the text). The events comprising E include transcription initiation, mRNA elongation and processing, nuclear export and mRNA decay, and cytoplasmic protein translation. The total system output—the amount of fluorescent reporter protein *O* produced in any cell *i*—depends on the product of *P*, *E*, and the duration of stimulation *ΔT*, with *P* itself dependent on α-factor concentrations. *P* and *E* are each split into an average component and a stochastic component: E = G + γ and P = L + λ. In each case the stochastic component is indicated by a Greek letter. The following equations describe the relationships among these quantities:

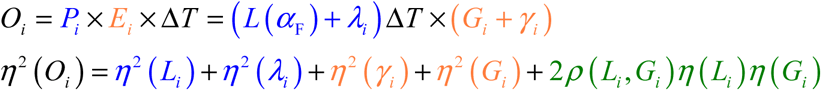 The function *η*^2^() is defined as the ratio of the variance of the indicated quantity to the square of the mean of the corresponding average component. So, for *O*, *G*, and *L*, *η*^2^() is equivalent to the square of the coefficient of variation (C.V.), but for the stochastic components *γ* and *λ*, the denominators are the squares of *G* and *L*, respectively.

#### Quantifying different contributions to cell-to-cell variation in cell signaling and response

Since DelbrÜck, non-genetic non-environmental cell-to-cell variation is sometimes attributed to statistical fluctuations in the output of biochemical processes, such as gene expression, that involve small numbers of protein components (Arkin et al., 1998; Delbrück, 1945; McAdams and Arkin, 1997). For example, Arkin and McAdams showed by modeling that stochastic fluctuations in gene expression could plausibly account for whether an infecting lambda phage lyses the cell or forms a lysogen (McAdams and Arkin, 1997). Such variability is often referred to as “noise” (Elowitz et al., 2002). This term can sometimes connote rapid fluctuations. However, other work reveals the importance of additional slowly-changing sources of variation in reducing coherence of population responses. In phage λ, incoming phage are more likely to lysogenize small cells than big ones, suggesting that one slow-changing source of variation (the cell growth and division cycle), rather than a fast-changing one ("noise" in the chemical reactions), causes the observed variability in the percentage of infecting phage that form lysogens (St-Pierre and Endy, 2008). Earlier work in *S. tymphimurium* showed that each individual bacteria retains its characteristic chemotactic behavior throughout its lifetime (Spudich and Koshland, 1976). Our work in the yeast pheromone response (Colman-Lerner et al., 2005, see below) revealed and quantified two slow-changing sources of variation, which we called P and G. Similarly, in mammalian cells abundance of particular apoptosis regulators determines the different timing of apoptosis in individual cells (Spencer et al. 2009). The abundance of these regulators in sibling cells is similar, and thus weakly heritable (Spencer et al., 2009). Such slow-changing cell-to-cell differences in protein abundance in cultured mammalian cells can, in some cases, predict drug response outcomes (Cohen et al., 2008).

In contrast, fast-changing fluctuations in protein concentration are unpredictable. For example, in the *lac* operon, variability in time to induction is caused by infrequent stochastic bursts in gene expression that arise from infrequent (in the order of once per generation) unbinding of the lacI repressor from its operators on the lac promoter (Choi et al., 2008).

Work by Elowitz et al. (2002), measured expression of two different colored fluorescent proteins driven by different instances of the same artificial LacI (lac repressor)-regulated promoter (Lutz and Bujard, 1997) in populations of clonal *E. coli*. This work defined two quantities: “intrinsic noise”, a measure of the extent to which the output of the two reporters did not correlate, and “extrinsic noise”, a measure of the remaining, correlated variation. Intrinsic noise, in most cases the smaller component, was presumed to arise from rapid-changing stochastic differences in the molecular events required for transcription and translation. Extrinsic noise was attributed to cell-to-cell fluctuations in the abundance of molecules such as regulatory proteins and polymerases. Both types of noise increased cell to cell variation in gene expression in the the cell population.

To dissect contributions to cell-to-cell variation to system output in the signaling arm of the yeast PRS, we used pairs of transcriptional reporters driven by different pairs of promoters (identical and non-identical, pheromone-inducible vs pheromone-insensitive). This experimental setup allowed us to separate cell-to-cell variation in molecular events upstream of the promoter (affecting a signal transmission or “pathway” subsystem, P) from those downstream (affecting a “gene expression” subsystem). It allowed us to further separate the contributions to variation in the gene expression subsystem caused by stochastic variation (γ) (Figure 1b) from that caused by preexisting differences in ability of cells to express proteins (G), and differences in signal transmitted by individual cells (P) (Figure 1c). We quantified variations in gene expression due to stochastic gene expression noise, γ, by comparing outputs in each cell of genes carrying α-factor-responsive promoters driving the YFP and CFP reporter genes (1b, drawn after Figure 1b in Colman-Lerner et al. 2005)). We then measure variation in pathway subsystem output (P) and expression capacity (E) in strains containing a pheromone responsive promoter driving YFP and a control promoter driving CFP reporter genes. (1c, drawn after 1c in Colman-Lerner et al. (2005)). Here, different Pathway subsystems (blue boxes) regulate the activity of the DNA-bound transcription factors, but the subsystem enabling expression of the reporter genes (red box) is the same. Variation in expression capacity, G, affects the correlated variation (the dispersion of points along the diagonal. Uncorrelated variation (the dispersion of points along the minor axis) is due to the stochastic gene expression noise, γ, and to cell-to-cell variations in the pathway subsystems for each promoter. Although this analytical framework recognized contributions to differences in transmitted signal (P) caused by pre-existing differences in the ability of cells to send signals (L) and stochastic differences in the ability of cells to send signals during the course of the experiment (λ), the experiments above do not allow us to distinguish them experimentally.

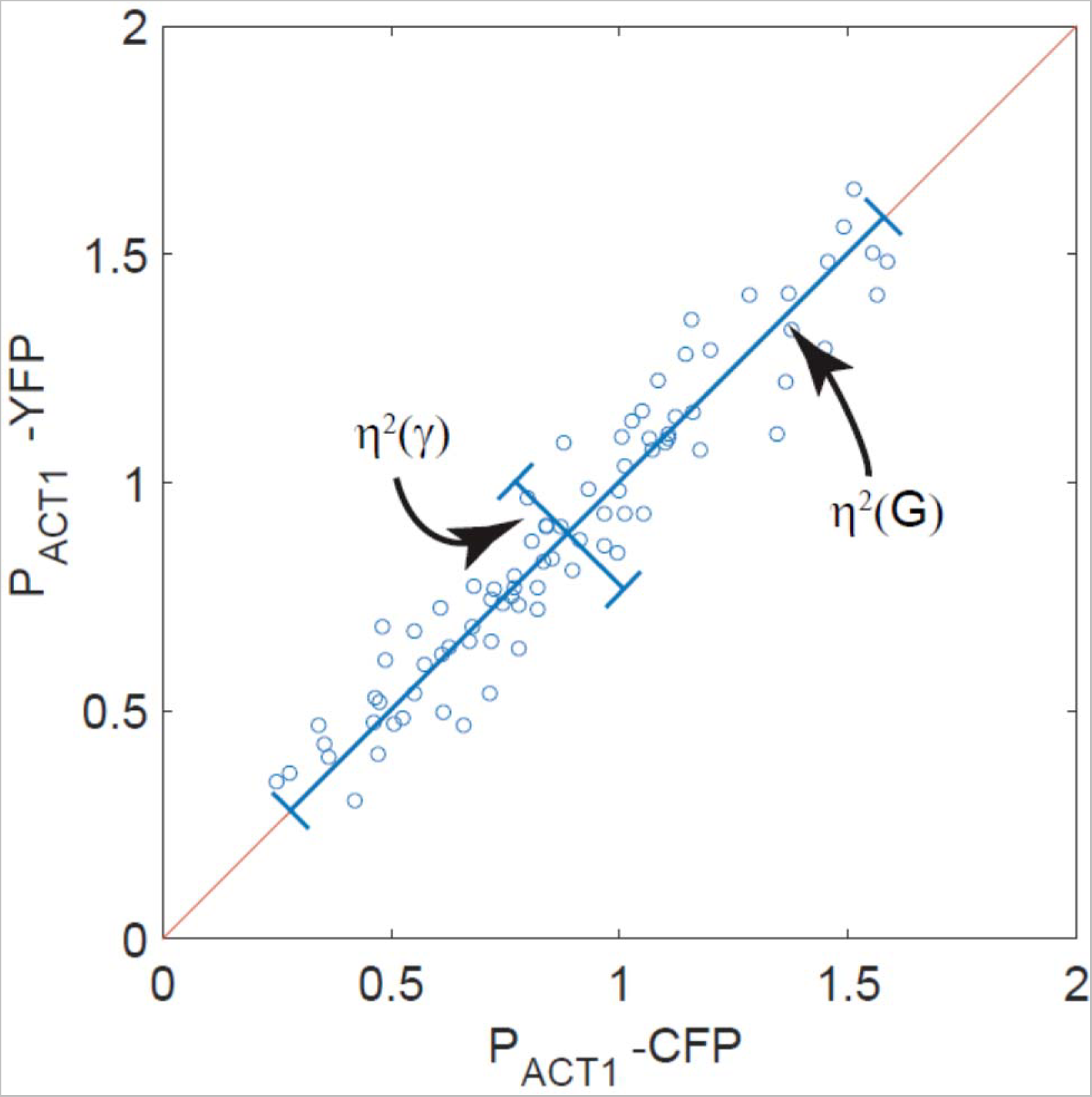
B. Measurement of gene expression capacity G and gene expression noise γ. In this illustrative example, measurement depends on quantification in each cell of YFP and CFP reporter genes each driven by a different instance of the same constitutive P_ACT1_ promoter. The spread of points along the correlation line shows cell-to-cell differences in the general ability of cells to express genes into proteins (η^2^(G)), while the spread of points across the correlation line corresponds to stochastic variation during the time of the experiment in the molecular processes needed for gene expression (η^2^(γ)).

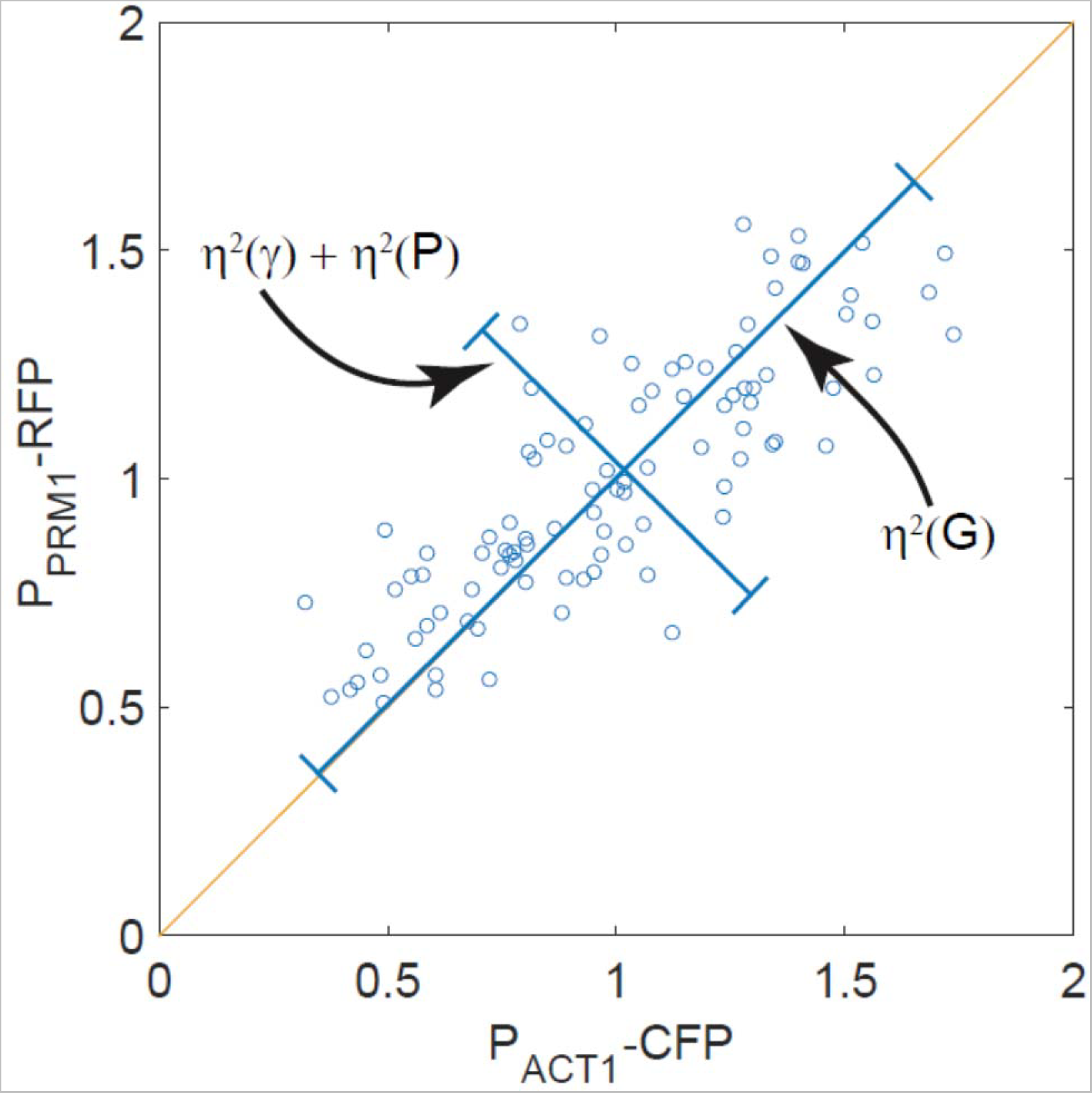
C. Measurement of variation in signal strength η^2^(*P*). Cell-to-cell variation in signal strength (P) and expression capacity (E, or G + γ) as measured in this work. Strains used in this figure carry a pheromone responsive promoter P_PRM1_ driving RFP and a second reporter in which a control, constitutive P_ACT1_ promoter drives CFP. In each cell, the Expression subsystem is the same, but the activity of the P_PRM1_ promoter depends on the accumulated signal transmitted to the promoter by the PRS. Again, expression of the two reporters is correlated (see the dispersion of points along the main diagonal), due to cell-to-cell differences in gene expression capacity G. Uncorrelated variation (the dispersion of points along the minor axis) is due to the combined effects of stochastic gene expression noise γ and to cell-to-cell variations in transmitted signal P for the inducible promoter P_PRM1_. Because there is only one instance of the P_PRM1_ promoter in these cells, and determination of γ for P_PRM1_ would require use of two different P_PRM1_ reporters as in 1b, experiments like the one shown here do not allow us to measure γ. They do however allow us to compute *η*^2^(*P*) + *η*^2^ (γ), and to use that as an estimate for *η*^2^ (*P*), for each cell (derivation in SI), since *η*^2^ (*γ*) is much smaller than η^2^(*P*). They cannot, however distinguish between the components of variation in P, i.e. variation in *L* (the pre-existing component of P,differences in the ability of cells to send signals), and the stochastic component, λ, due for example to differences caused by differences in molecular events needed for signaling during the course of the experiment.

We quantified total cell-to-cell variability using the normalized variance η^2^, the variance squared over the mean squared, σ^2^/μ^2^ (for G, P, E, and L, this is equivalent to the square of the coefficient of variation or CV). We also quantified three components of η^2^_total_:“Cell-to-cell variability in gene expression capacity, G”, η^2^(G) (the overall capacity of a cell to transcribe genes into mRNAs and translate those mRNAs into proteins), “stochastic variability in gene expression” or “gene expression noise”, η^2^(γ) (which corresponded to “intrinsic noise”), and “cell-to-cell variability in pathway subsystem output”, η^2^(P). Both η^2^(G) and η^2^(γ) reduced the coherence in population gene expression responses. In these experiments, we found the contribution of η^2^(γ) or intrinsic noise was very small. Even when we eliminated variation due to cell cycle position, most variation was due to differences in P and in G (Colman-Lerner et al., 2005). As mentioned above, η^2^(P) is the sum of two components, which we could not separate experimentally: η^2^(L), and η^2^(λ).

Three lines of evidence show that η^2^(P) (here called cell-to-cell variability in transmitted signal, or signaling variation) is under active control. First, pathway subsystem output P correlates negatively with gene expression capacity G, defining a compensatory mechanism that reduces variation in transmitted signal. Second, both PRS MAPKs, Kss1 and Fus3, regulate η^2^(P). At low pheromone inputs, *Δkss1* cells showed decreased η^2^(P) while at high pheromone inputs *Δfus3* cells showed increased η^2^(P) indicating that the products of these genes control variation in transmitted signal (Colman-Lerner et al.). Third, the dose-response relation for gene expression output and for several intermediate signaling steps matches the dose-response relation for fractional receptor occupancy (Brent). This <u>Do</u>se-<u>R</u>esponse <u>A</u>lignment (or DoRA), is a “systems level” behavior that improves the fidelity of information transmission by making downstream responses more distinguishable, and also, under some circumstances, by reducing amplification of stochastic noise n^2^(λ) during signal transmission (Yu et al., 2008). DoRA operates in many other signaling systems including insulin, (Cuatrecasas, 1971), angiotensin II (Lin and Goodfriend, 1970) and EGF (Knauer et al., 1984). The fact that in the PRS and other systems DoRA is present after long exposure to ligand and in the face of significant changes in the numbers of signaling molecules (Knauer et al., 1984; Thomson et al., 2011) suggests that DoRA is actively maintained. In the PRS, maintenance of DoRA requires the action of a particular negative feedback (Yu et al., 2008).

We realized that mutations affecting mechanisms that reduced variation in different components of η^2^(P) should have different effects on the responses of cell populations. Interference with mechanisms that reduced either component of pathway variation (η^2^(L) or η^2^(λ)) would increase cell-to-cell variability in transmitted signal and thus decrease the coherence of the response of *cell populations*. However, interference with mechanisms that suppressed differences in signal transmission due to the stochastic component η^2^(λ) (caused for example by differences in molecular collisions or other events during the course of an experiment) would also reduce the precision in the response of *individual cells*. Here, we undertook a comprehensive genetic screen for genes that regulated different components of cell-to-cell variability in the yeast *S. cerevisiae* pheromone signaling and response. This work identified a previously unknown and unexpected role for cytoplasmic microtubules in reducing cell-to-cell variation in transmitted signal and increasing coherence of the population response.

## Results

### Construction of a whole-genome collection of viable single gene deletions permitting screens for variation in signal transmission and response

To introduce the necessary reporters and alleles for the screen, we extended methods of genetic cross and segregant selection described previously (Tong et al., 2004). We constructed a MAT α strain, SGA88, with additional antibiotic resistance markers, auxotrophies, and reporter genes (described in SI). We used SGA88 to cross to an otherwise-isogenic collection of MAT α strains, each one of which carried a deletion in each non-essential gene in the yeast genome (about 4,100 strains) (Giaever et al., 2002, Giaever and Nislow, 2014). We then sporulated this diploid strain collection, and selected MATa haploids that carried the deletion plus our necessary reporters and alleles, resulting in a whole-genome collection suitable for the cell to cell variation screen (Figure 2a, Methods, and SI). Importantly, to ensure that all members of this collection were clonal, derived from single founder spores, we used cultures derived from single colonies streaked from small colonies that had germinated on the 8-way selective medium used to isolate MATa haploids with the appropriate genotype. Our generation of this strain collection from single colonies stands in contrast to the usual practice of growing cultures from patches derived from thousands of germinated spores (derived from thousands of independent meioses) (Ayer et al., 2012; Jonikas et al., 2009; Neklesa and Davis, 2009; Wolinski et al., 2009). Because these cultures were clonal, we hoped to ensure genetic homogeneity. By using cultures made from independent, newly isolated small colonies in particular, we ensured that those phenotypes we observed consistently in initial screens derived from the deletions and not from unlinked second-site mutations, either pre-existing in the founder haploid deletion collection or that arose during meiosis and sporulation, that suppressed a growth defect caused by the deletion (Hittinger and Carroll, 2007). Moreover, by relying on cultures derived from freshly germinated spores generated by new meioses, we circumvented possible complications of aneuploidy (Hughes et al. 2000) and same-sex diploid formation (Giaever and Nislow 2014) that might have affected members of the starting deletion collection.

**Figure 2.**
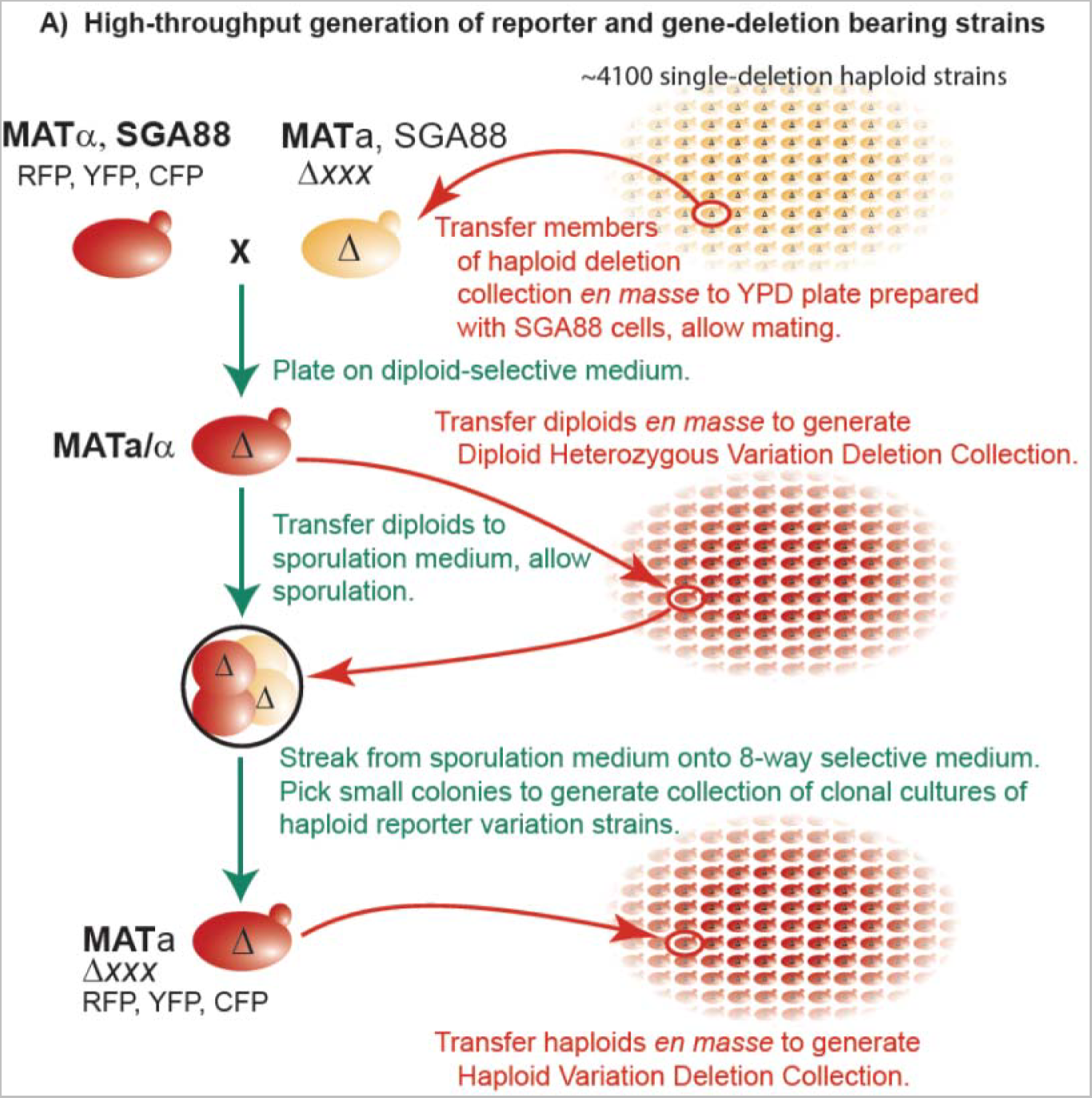
Strain modification protocol and generation of haploid reporter collection. **A. High throughput generation of reporter and gene-deletion bearing strains.** We crossed the BY4742 derivative SGA88b *(MATα Δcan1::P_MFA1_-LEU2 Δbar1-orf::P_PRM1_-CFP–HIS3 Δbar1-promoter::ura3-terminator Δlyp1::P_ACT1_-YFP–URA3 Δprm1::P_PRM1_-RFP–NAT(MX4) cdc28-F88A–hph(HygB^r^)(MX4))* to strains in yeast *MATa* haploid deletion collection *(Axxx::G418 (MX6))where* xxx is the deleted yeast gene) using 384 pinning tools (see SI),. We sporulated the resulting diploids, and selected spores of the desired genetic makeup: *MATα Δxxx Δcan1::P_MFA1_-LEU2 Δbar1-orf::P_PRM1_-CFP–HIS3 Δbar1-promoter::ura3-terminator Δlyp1::P_ACT1_-YFP–URA3 Δprm1::P_PRM1_-RFP—NAT(MX4) cdc28-F88A–(hph(HygB^r^)(MX4))*, grew these into single colonies and re-streaked these to obtain the haploid collection used in this screen.

Figure 2b shows the key markers in strains in the collection. All strains carried two fluorescent protein reporters controlled by *P*_*PRM1*_, the promoter of the pheromone inducible gene *PRM1* (*P*_*PRM1*_-*mRFP* and *P*_*PRM1*_-*CFP*) (Colman-Lerner et al., 2005) and a third fluorescent protein reporter controlled by *P_ACT1_*, the promoter of the housekeeping gene *ACT1* (Colman-Lerner et al., 2005) (Figure 2B). They also carried a deletion of the *BAR1* protease gene, to ensure consistent extracellular pheromone concentrations, and the *cdc28*-*as2* allele (F88A) in the *CDC28* locus. The variant of Cdc28 encoded by *cdc28*-*as2* renders the single cyclin-dependent cell cycle kinase in yeast sensitive to inhibition by the ATP-analog 1NM-PP1 (Bishop et al., 2000). Inhibiting Cdc28 allowed us to minimize cell-cycle-dependent variation in pheromone response and to block cell division (and thus dilution of fluorescent protein into new daughter cells) (Colman-Lerner et al., 2005).

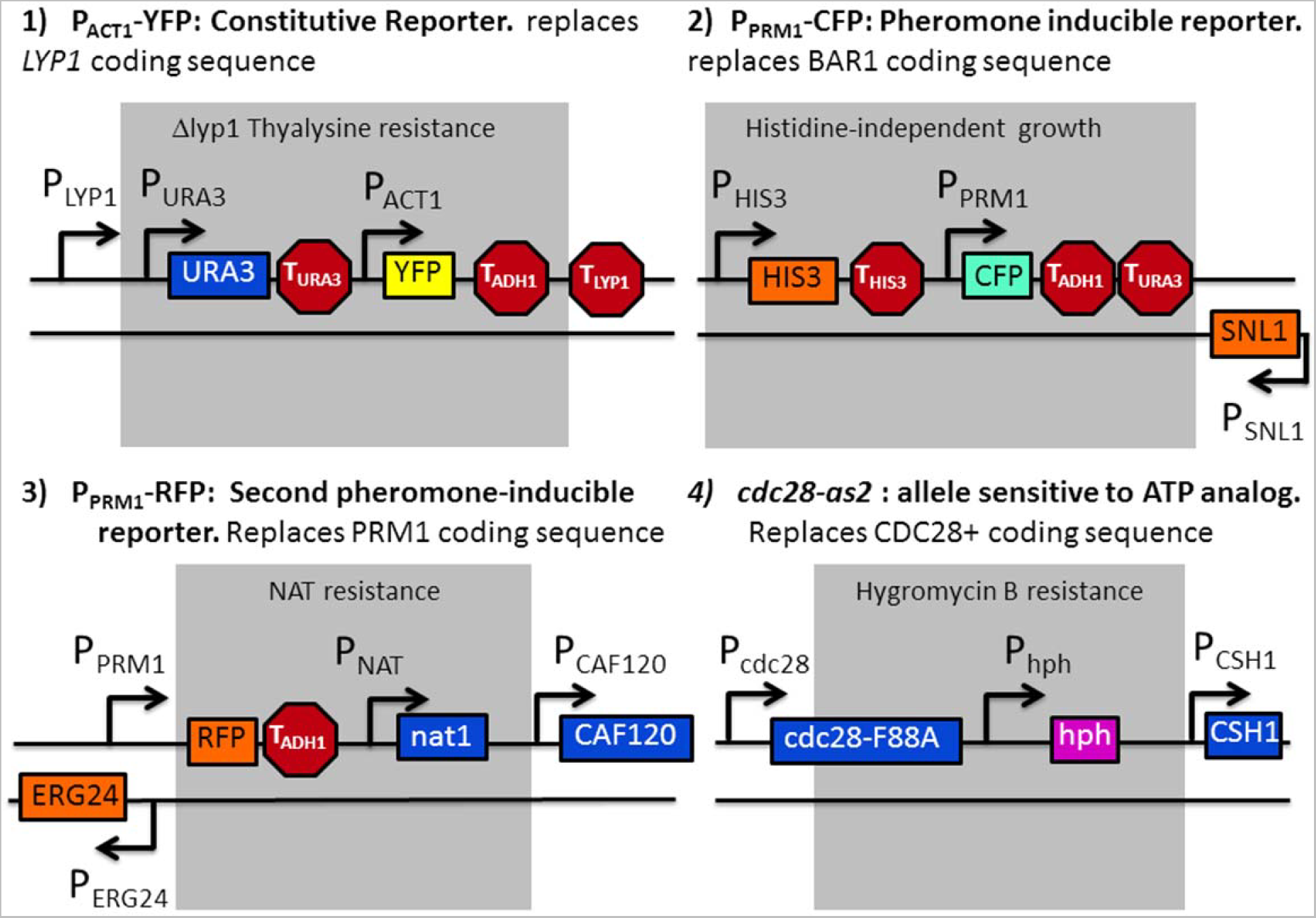
B. Key markers in strains in the reporter collection. **(1)** P_ACT1_-YFP constitutive reporter. Replaces LYP1 coding sequence. LYP1 deletion allows selection for diploids. **(2)** P_PRM11_-CFP inducible reporter, replaces BAR1 coding sequence. (3) P_PRM11_-RFP Second inducible reporter, replaces PRM1 coding sequence. (4) cdc28-as2 allele installed in place of CDC28+ coding sequence. confers sensitivity to the ATP analogue 1-NM-PP1, causing cell cycle arrest.

To prepare to screen these strains for variation in signal and response, we measured mRFP and YFP signal in the reference strain SGA85 by flow cytometry after pheromone induction. (Methods, Figure 3a and 3b). We approximated the cell-to-cell variability in signal strength η^2^(P) as the portion of the variation in mRFP signal that did not correlate with the YFP signal (calculated as the variance of the differences in the normalized expression of two non-identical reporters (Figure 1c, see SI for derivation)) (Colman-Lerner et al., 2005). We used this method to extend our previous characterization of the PRS. We did so by inducing system operation with a broad range of different pheromone concentrations (0.1nM to 30 nM). As expected, system output increased with dose (EC50 of 0.8 nM) (Figure S1). In contrast, signaling variation η^2^(P) decreased monotonically with increasing pheromone input (Figure S1), consistent with our previous microscopic (image cytometric) measurements (Colman-Lerner et al., 2005). As a second test of our flow cytometric screening methods, we verified that, as previously observed, derivatives of the reference strain lacking Fus3 showed increased η^2^(P) at high pheromone inputs, while derivatives lacking Kss1 showed reduced η^2^(P) at low inputs (Figure S2).

**Figure 3.**
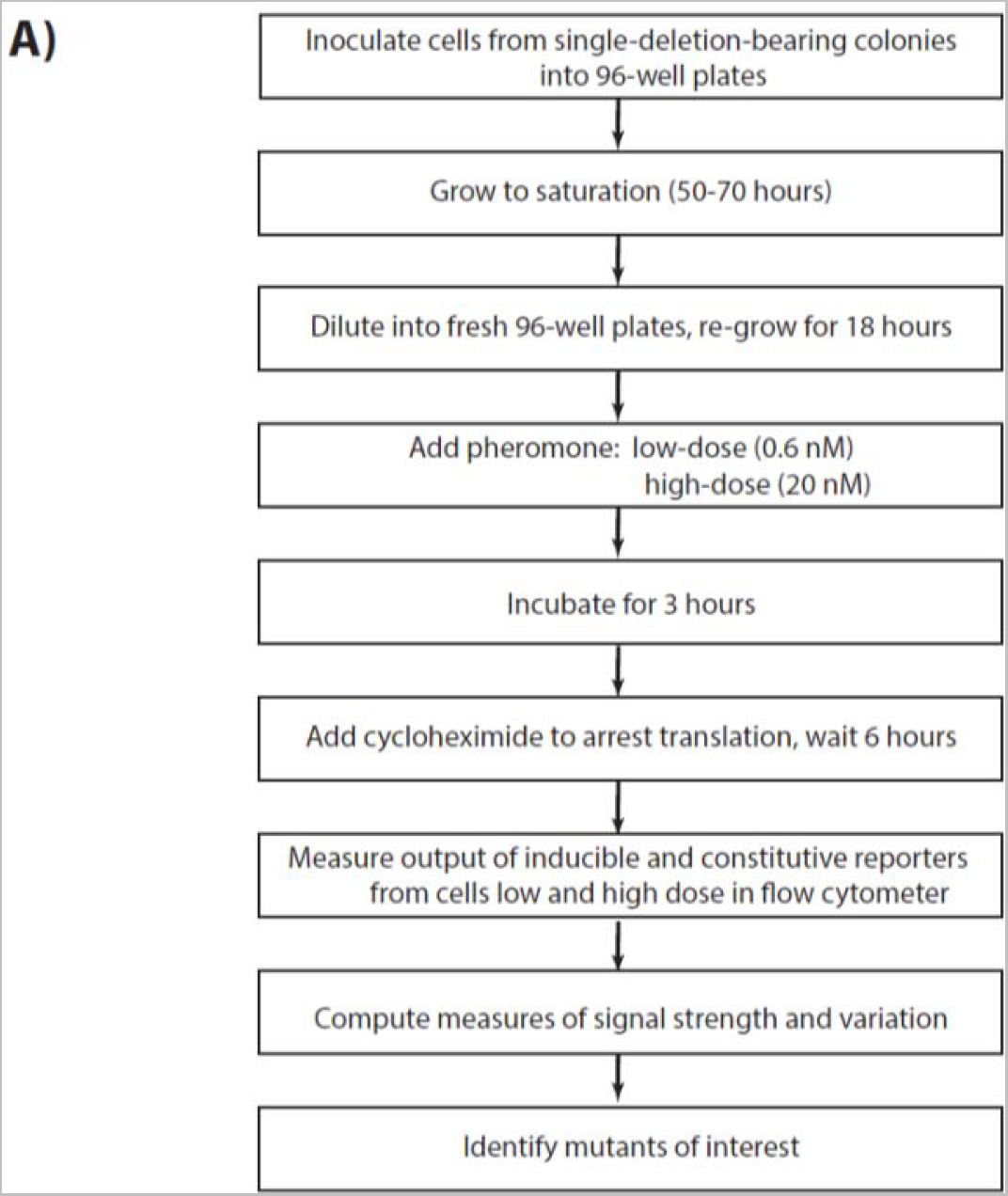
High throughput genetic screen for mutants in signal variation η^2^(P), signal strength P, or system output O. **A. Flow chart showing steps in screening for strains altered in pathway variation or output.**

### Large-scale screen identifies genes whose products affected cell-to-cell variation in transmitted signal

We used these methods to carry out a high throughput screen for genes that, when deleted, altered η^2^(P). For the primary screen, we screened 1141 strains from the collection (996 randomly selected, together with an additional 145 which each bore a deletion in a nonessential kinase or phosphatase (Supplementary information, Table S1), for mutants with altered values of pathway output and/or η^2^(P). Screened strains corresponded to more than 1/4 of the nonessential yeast genes. Figure 3a describes the steps we followed. We grew one instance of each haploid deletion strain, plus 50 cultures of the reference strain (as controls), in log phase (less than 3 × 10^6^ cells/ ml) for at least 14 hours, and then exposed them for 3 hours to two different (0.6 nM and 20 nM) pheromone concentrations in the presence of 10μ 1-NM-PP1. We then added 100 μg/ml cycloheximide to inhibit protein synthesis and allowed existing translated fluorescent protein molecules to mature (Colman-Lerner et al., 2005; Gordon et al., 2007) (Figure 3a), and took other steps to ensure accuracy and consistency of measurement described in SI. We measured fluorescent signal by flow cytometry as shown in Figure 3b. From these measurements, we calculated values for 5 variables of interest shown in Table 1. These were: Average output (O), cell-to-cell variability in output, η^2^(O), median output of the P_ACT1_ constitutive reporter (an estimate of G), cell-to-cell variability in this output, η^2^(G), and cell-to-cell variability in transmitted signal, η^2^(P), at each of the two pheromone doses.

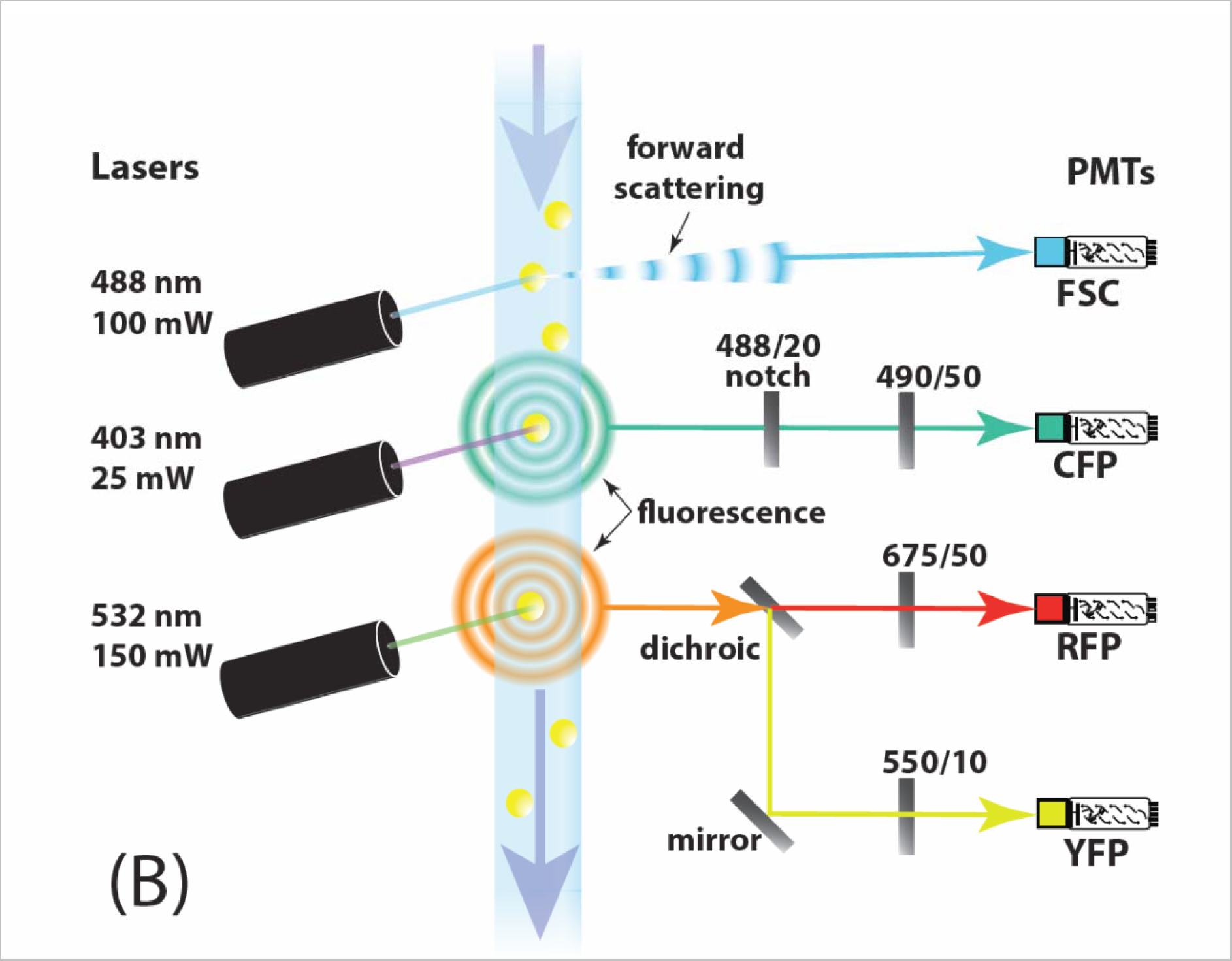
B. Light path enabling three-color flow cytometric genetic screen. Forward-scattering of 488 nm laser light indicates the presence of cells (yellow), while fluorescence in three separate wavelength bands indicates the quantity of cyan, red, and yellow fluorescent reporter proteins within cells. Photomultiplier tubes detected scattering and fluorescence intensities and recorded these in channels correspondingly labeled FSC, CFP, RFP, and YFP. Barrier bandpass filters, each labeled above with the center wavelength of the passband and the bandwidth in nanometers, ensured that excess laser light did not enter the PMTs. An additional bandstop filter ("notch"), labeled with the center wavelength and bandwidth of the stop band, ensured that light from the 488 nm laser did not enter the CFP PMT.

**Table 1.** Variables measured in isogenic cell populations

legend. $\sigma^2$ is variance, $\mu$ is average, $\langle B \rangle$ means the average of B
| Table 1. Variables measured in isogenic cell populations |  |  |
| --- | --- | --- |
| Variable name | Short Expression | Calculated as |
| Median pheromone response system (PRS) output | **O**, $\langle P_{PRM1}\text{-mRFP} \rangle$ or $\langle P_{PRM1}\text{-mCherry} \rangle$ | median(inducible RFP) |
| Median constitutive or control output | **G**, $\langle P_{ACT1}\text{-XFP} \rangle$ or $\langle P_{BMH2}\text{-XFP} \rangle$ | median(constitutive XFP) |
| Cell-to-cell variability in PRS output | $\eta^2(\mathbf{O})$, $\eta^2(P_{PRM1}\text{-mRFP})$ or $\eta^2(\text{mCherry})$ | $\sigma_{XFP}^2 / \mu_{XFP}^2$ |
| Cell-to-cell variability in constitutive or control output (representing general gene expression capacity) | $\eta^2(\mathbf{G})$, $\eta^2(P_{ACT1}\text{-YFP})$ or $\eta^2(P_{BMH2}\text{-YFP})$ | $\sigma_{YFP}^2 / \mu_{YFP}^2$ |
| Cell-to-cell variability in signal transmission | $\eta^2(\mathbf{P})$, or $\eta^2(L+\lambda)$ | $\sigma^2 (mRFP_i / \langle mRFP \rangle - YFP_i / \langle YFP \rangle)$ |
| Signal strength | **P** | **O/G** |

We selected for follow up studies all gene deletions that exhibited high or low median pheromone system output (O) (Figure 4a) or significantly higher or significantly lower η^2^(P) than the reference strain at the low dose (Figure 4b), and strains that showed higher η^2^(P) than the reference strain at the high dose (Figure 4c). Figure 4d shows an overview of signaling variation vs. output, at the two pheromone doses, for all measured strains. To perform follow-up, and to again avoid possible complications from second-site suppressor mutations, we isolated multiple instances of independent haploid clones of deletion strains derived from the diploid parent strain. We then rescreened these independent isolates (see SI for a complete description of primary and secondary screens) by flow cytometry as above. We treated measurements from each of three or four isolates of each deletion (each a clone of isogenic cells descending from a different isolated haploid spores) as separate entries. These experiments identified 50 strains (Table 2) bearing deletions in candidate variation genes that showed variation in O or η^2^(P) beyond the thresholds (Figure 4a, 4b, and 4c). In 44 of these 50 mutants, we tested for and indeed reproduced altered O and/or η^2^(P) (see SI and Table S6) by microscope-based quantification of accumulated fluorescent signal in groups of several hundred single cells. In these verification experiments, we also quantified PRS gene expression noise η^2^(γ) using the two reporters (CFP and mRFP) driven by *P_PRM1_*. The tested mutants showed values of η^2^(γ) that were typical of the reference strain. The only significant differences were in O, η^2^(O), and η^2^(P).

**Figure 4.**
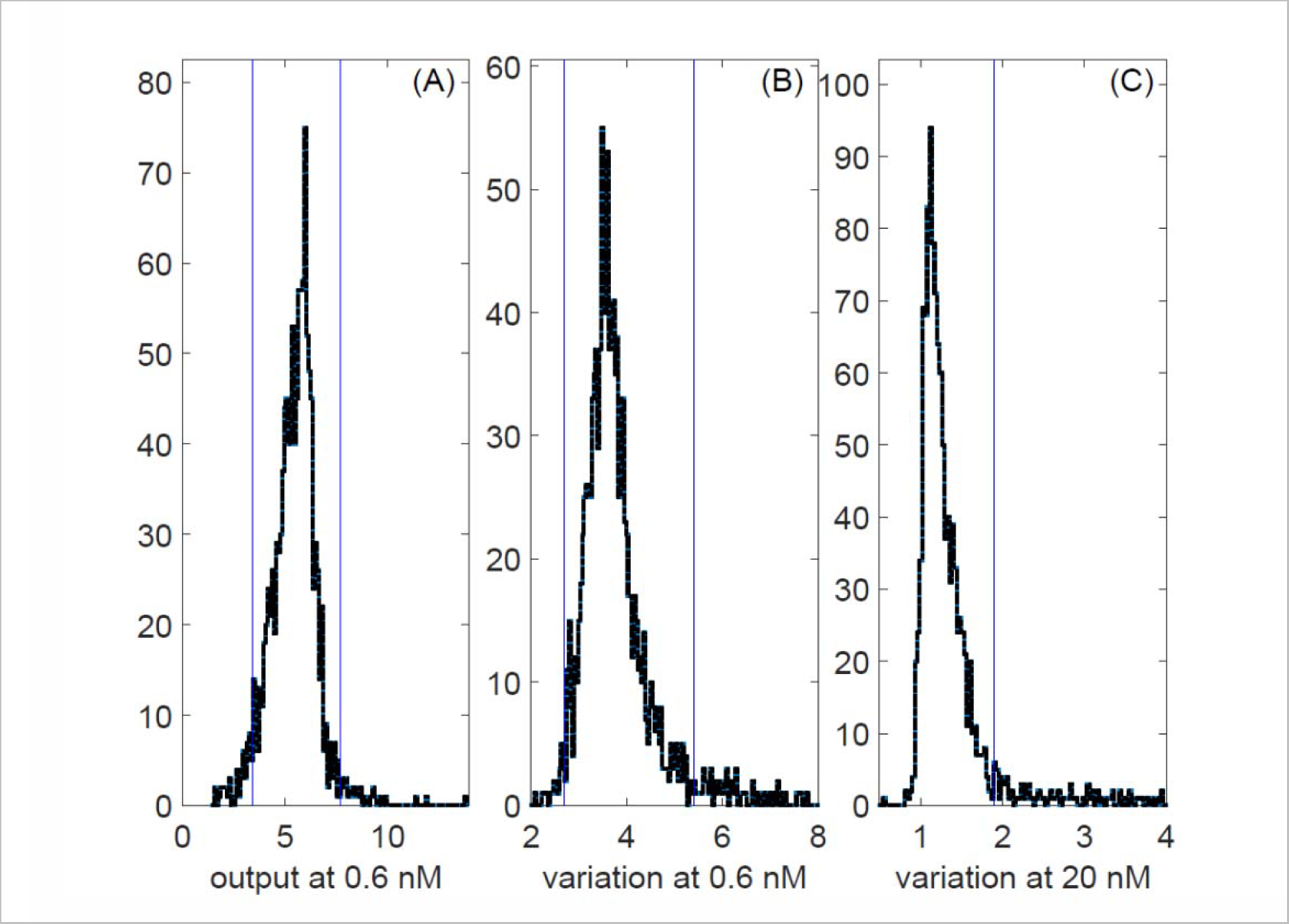
A-C. Selection of mutants for follow up studies. Plots show distributions of values for 991 randomly selected non-essential deletion strains, and 102 additional strains with deletions of a non-essential kinase or phosphatase, and 2 wild-type strains. Values were derived from flow cytometry data obtained after 3h of stimulation with pheromone. Blue vertical bars indicate the thresholds used to select mutants for secondary screens (see SI). **A.** PRS output, O (median mRFP signal from pheromone inducible reporter gene), in 0.6 nM pheromone. **B.** Estimated signal variation η^2^(P) in 0.6 nM pheromone **C.** Estimated signal variation η^2^(P) in 20 nM pheromone.

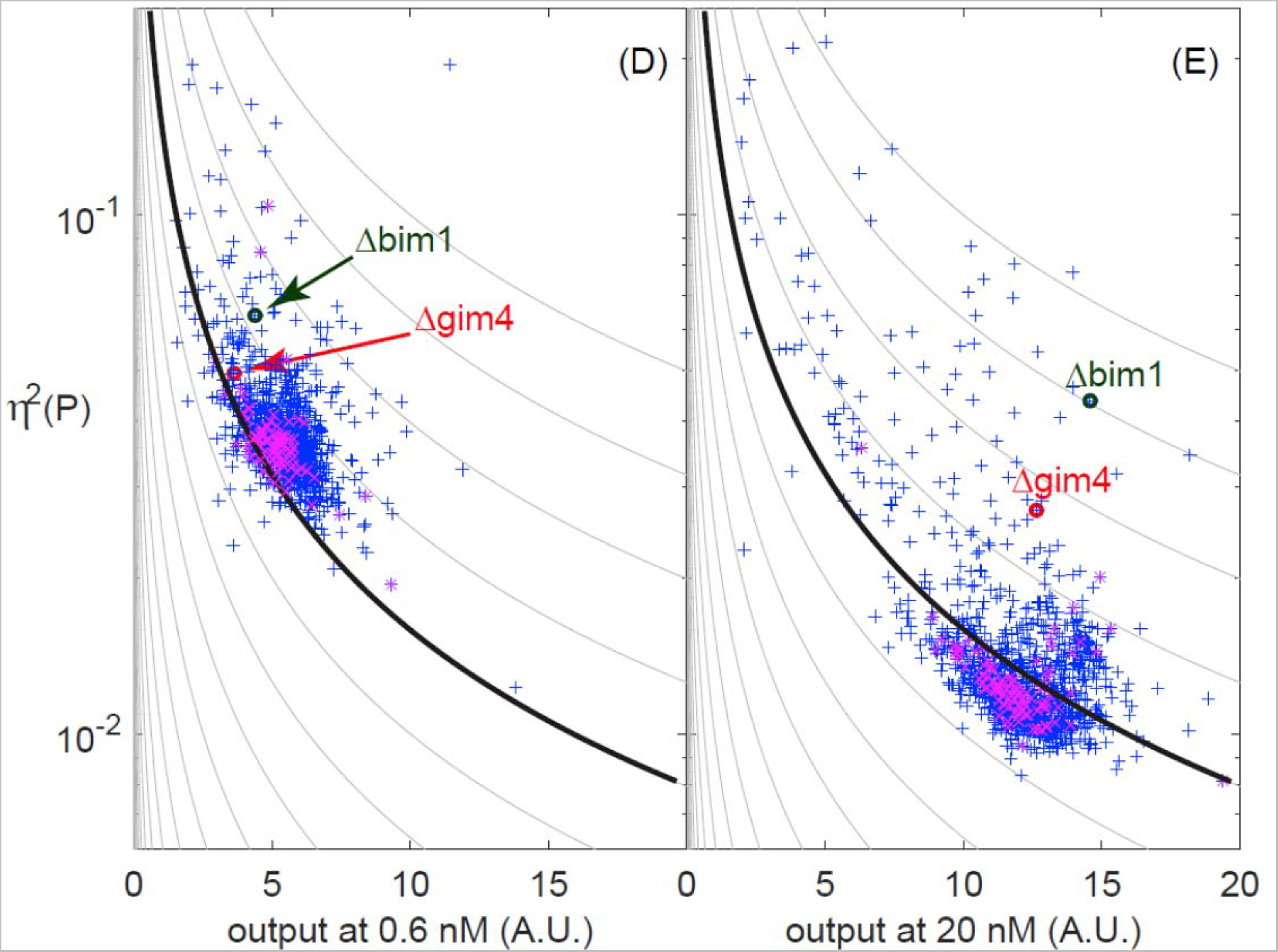
D-E. Signal variation versus output for all 1093 strains screened. Plots show an estimate of η^2^(P) vs. PRS output (O) for the same data set displayed in Figures 4A-C. Purple x’s are WT cells. At 20 nM, *Δbim1* and *Δgim4* show significantly less population coherence than WT. These strains were stimulated for 3 h with (D) 0.6 nM and (E) 20 nM pheromone. The contour lines show the expected dependence of variation on output for outputs proportional to a Poisson random variable (lower noise at higher outputs), with proportionality constants logarithmically spaced from 10^-5^ to 1. The WT swarm lies below the 0.158 contour at 20 nM but above it at 0.6 nM, indicating that variation at the low dose is higher than expected from the same Poisson processes that take place at 20nM. See Supplemental file TableS1.xlsx for a list of all 1095 strains and their corresponding output and variation values.

**Table 2.** Genes found in the screen

| Gene name | Screen | Important for | Description |
| --- | --- | --- | --- |
| *ARG82* | UnB | AA metabolism | Inositol polyphosphate multikinase |
| *EVR46* | UnB | Cargo transport | ER vesicle protein, component of COPII complex; required for membrane fusion |
| *HIS1* | UnB | AA metabolism | ATP phosphoribosyltransferase |
| *UGA1* | UnB | AA metabolism | Gamma-aminobutyrate (GABA) transaminase |
| *SLA1* | UnB | Actin binding | Cytoskeletal protein binding protein; required for assembly of the cortical actin cytoskeleton |
| *SAP155* | K&P | Cell cycle | Protein required for function of the Sit4 protein phosphatase |
| *YER068C-A* | UnB | Dubious open reading frame | Dubious open reading frame/ overlaps with ARG5, ARG6 Acetylglutamate kinase and N-acetyl-gamma-glutamyl-phosphate reductase |
| *YIL032C* | UnB | Dubious open reading frame | Dubious open reading frame / next to BCY1 |
| *ERG3* | UnB | Ergosterol biosynthesis | C-5 sterol desaturase |
| *GAL83* | K&P | Glucose repression | One of three possible beta-subunits of the Snf1 kinase complex |
| *GUP1* | UnB | Glycerol metabolism, protein folding | Plasma membrane protein involved in remodeling GPI anchors |
| *PPZ1* | K&P | Ion homeostasis | Serine/threonine protein phosphatase Z, isoform of Ppz2p; involved in regulation of potassium transport, which affects osmotic stability, cell cycle progression, and halotolerance |
| *FUS1* | UnB | Mating | Membrane protein localized to the shmoo tip |
| *AAT2* | UnB | Metabolism | Cytosolic aspartate aminotransferase involved in nitrogen metabolism |
| *PTC6* | K&P | Metabolism | Mitochondrial type 2C protein phosphatase (PP2C) |
| *GIM4* | UnB | Microtubule chaperone/<br>Protein folding | Subunit of the heterohexameric cochaperone prefoldin complex |
| *PAC10*<br>*GIM2* | UnB | Microtubule chaperone/<br>Protein folding | Subunit of the heterohexameric cochaperone prefoldin complex |
| *BIM1* | UnB | Microtubule end binding | Microtubule plus end-binding protein |
| *MSH1* | UnB | Mitochondrial homeostasis | E. coli MutS homolog, binds DNA mismatches, required for mitochondrial function |
| *BUB1* | K&P | mitosis | Protein kinase required for cell cycle checkpoint, delays entry into anaphase until kinetochores bound by opposing microtubules |
| *ELM1* | K&P | morphogenesis | Serine/threonine protein kinase that regulates cellular morphogenesis |
| *HSL1* | K&P | morphogenesis | Nim1-related protein kinase; regulates the morphogenesis and septin checkpoints |
| *NUP60* | UnB | nuclear transport | FG-nucleoporin component of central core of the nuclear pore complex |
| *SXM1* | UnB | nuclear transport | Nuclear transport factor (karyopherin) |
| *CBR1* | UnB | respiration | Microsomal cytochrome $\beta$ reductase |
| *RTC3* | UnB | RNA metabolism | Protein of unknown function involved in RNA metabolism |
| *CKA1* | UnB | signaling | Alpha catalytic subunit of casein kinase 2 (CK2) |
| *CKB1* | UnB | signaling | Beta regulatory subunit of casein kinase 2 (CK2) |
| *CKB2* | K&P | signaling | Beta' regulatory subunit of casein kinase 2 (CK2) |
| *FUS3* | K&P | signaling | Mitogen-activated serine/threonine protein kinase (MAPK), part of PRS |
| *HOG1* | K&P | signaling | Mitogen-activated protein kinase involved in High Osmolarity (HOG) pathway |
| *KSS1* | K&P | signaling | Mitogen-activated protein kinase (MAPK); functions in PRS and signal transduction pathways that control filamentous growth and pheromone response |
| *PBS2* | K&P | signaling | MAP kinase kinase of the HOG signaling pathway |
| *SSK2* | K&P | signaling | MAP kinase kinase kinase of HOG1 signaling pathway |
| *FAR1* | UnB | signaling/cell cycle/polarization | CDK inhibitor, nuclear anchor, recruited by Ste18-Ste4 at polarity patch |
| *SIP1* | UnB | signaling/glucose repression | Alternate beta-subunit of the Snf1 protein kinase complex |
| *KAR4* | UnB | signaling/mating | Transcription factor required for activation of some pheromone responsive genes |
| *STE50* | UnB | signaling/mating | Adaptor protein, in PRS helps connect Ste20 MAPKKKK to Ste11 MAPKKK |
| *SKY1* | K&P | splicing | SR protein kinase (SRPK); varied functions, regulates proteins involved in mRNA metabolism and cation homeostasis, helps some LexA fusion proteins bind operator |
| *KIN3* | K&P | stress | Nonessential serine/threonine protein kinase; possible role in DNA damage response |
| *OCA1* | K&P | stress | Protein tyrosine phosphatase; required for cell cycle arrest in response to oxidative damage of DNA |
| *CTK1* | K&P | transcription regulation | Catalytic (alpha) subunit of C-terminal domain kinase I (CTDK-I); phosphorylates RNA pol II |
| *DEP1* | UnB | transcription regulation | Component of the Rpd3L histone deacetylase complex, variously needed for activation and repression, regulates DNA replication origin timing |
| *SUM1* | UnB | transcription regulation | Transcriptional repressor that regulates middle-sporulation genes; required for mitotic repression of middle sporulation-specific genes; also acts as general replication initiation factor; involved in telomere maintenance, chromatin silencing |
| *SWI5* | UnB | transcription regulation | Part of Mediator and Swi/Snf nucleosome remodeling complexes |
| *UME6* | UnB | transcription regulation | Meiotic transcription regulator, DNA binding, recruits variously Sin3/ Rpd3 repressor (HDAC) and Ime1 activator. |
| *RPL12A* | UnB | Translation | Ribosomal 60S subunit protein L12A |
| *RPL19B* | UnB | Translation | Ribosomal 60S subunit protein L19B |
| *RPL34A* | UnB | Translation | Ribosomal 60S subunit protein L34A |
| *ECM15* | UnB | unknown | Possibly tetrameric, nonessential protein, unknown function |
| *VPS64* | UnB | Vacuole metabolism | Required for cytoplasmic proteins to enter vacuole |

The 50 genes from the 886 strains in the unbiased (UnB) screen and the 145 strains in the nonessential kinase and phosphatase screen (K&P) screen that showed altered PRS system output (O), low or high cell-to-cell variability in transmitted signal ( $\eta^2(P)$ ) at low pheromone (0.6 nM), or high variability in ( $\eta^2(P)$ ) at high (20nM). Table shows gene name, screen from which it was selected, overall functional class, and a brief description of its molecular role or activity. ERV46 is out of place in this table.

### Mutant genes define different axes of quantitative system behavior

To gain insight into the different phenotypes caused by these gene deletions, we clustered the mutant strains based on the 5 variables we had measured by flow cytometry at low and high doses (Table 1 and Table S2). We treated as separate entries the measurements from each of the three or four isolates of each gene deletion included in our secondary screens (each representing a clone of isogenic cells descending from a different isolated haploid spore, see SI) and the 19 instances of measurements for the reference strain. To generate the clusters, we used an uncentered Pearson correlation and average linkage method. All 19 cultures of the reference strain grouped together in one cluster (cluster I), along with the replicas from the *Δfus*1 deletion, which showed a weak phenotype (Figure 5).

**Figure 5.**
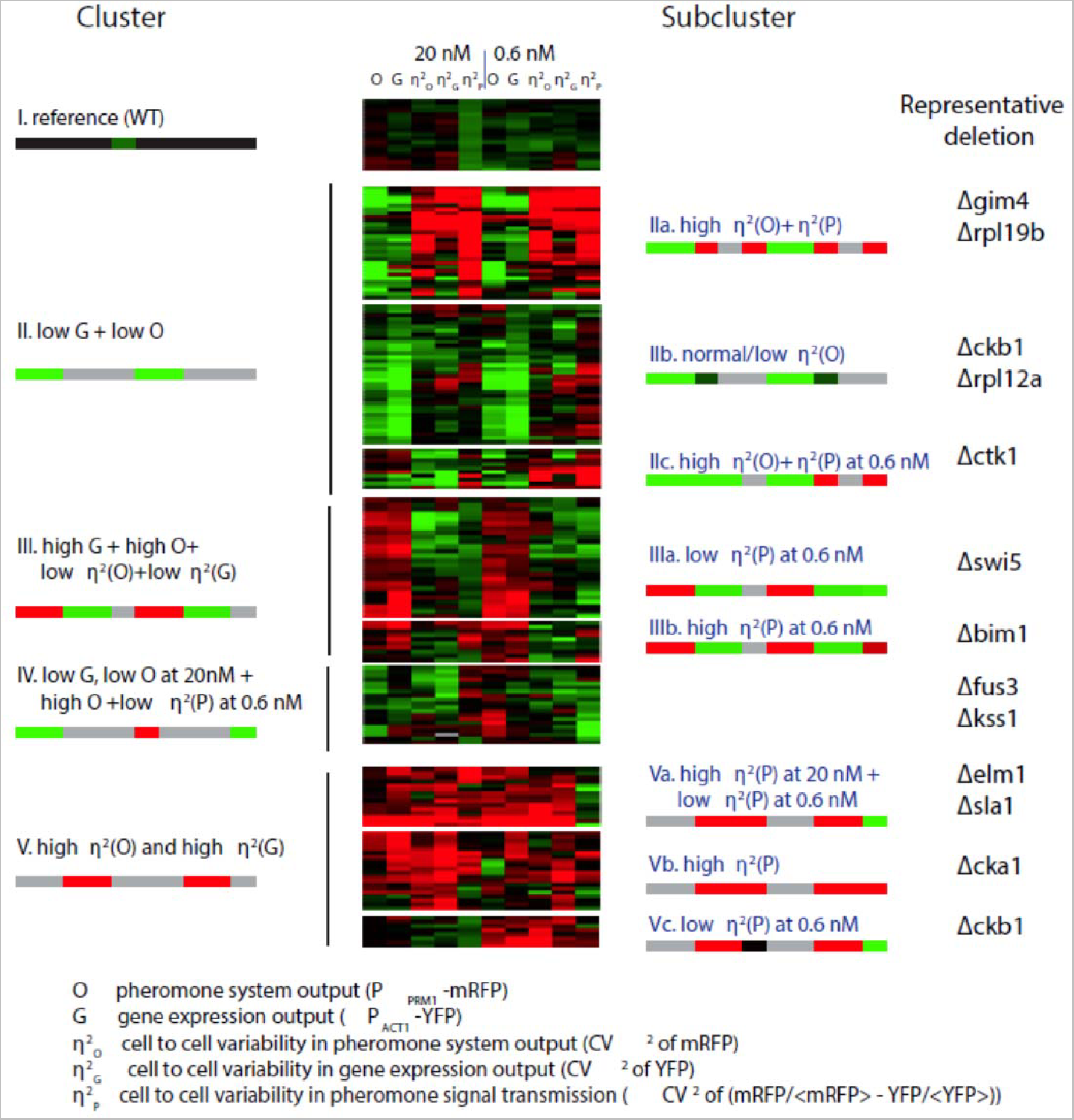
Cluster analysis of 50 genes identified as affecting variation and or pheromone response output. Hierarchical clustering of values derived from flow cytometry measurements (detailed in Methods and SI) from 193 strains (19 replicates for reference strain SGA85, 4 independent segregants each for 17 deletions from the kinases or phosphatase set and 3 independent segregants each for 37 deletions from the unbiased set). We used the Pearson correlation metric to assess distance between strains and the average linkage method to form clusters. Before clustering, we first log transformed the data, then median centered each row (each strain). Each strain had the following 10 measurements (5 after induction with 20 nM pheromone and 5 after induction with 0.6 nM pheromone): O (pheromone system output), G (gene expression output), η^2^(O) (cell-to-cell variability of the pheromone system output, the CV^2^ of the mRFP signal), η^2^(G) (cell-to-cell variability of the gene expression output, the CV^2^ of the YFP signal) and η^2^(P) (cell-to-cell variability in pheromone signal transmission, *σ*^2^ (*mRFPj_i_*/ < *mRFP* > − *YFP_i_* < *YFP* >). The panel shows these values as a “heat map”, from red (higher than the median) to green (lower than the median) through black (equal to the median). There were five main clusters (left, labeled I though V) and 1 to 3 subclusters (right). The signature pattern for each cluster or subcluster is represented with a color bar with 10 blocks, one for each measurement (grey indicates that that the measurement may take any value). Rightmost column shows representative deletion strains for each subcluster. Table S2 lists the data before transformation and Table S3 lists the clustered, log transformed and median centered dataset. Table S6 shows microscope data that complement these flow cytometer data for 44 of these 50 mutant strains selected for clustering.

Analysis of data from the deletion strains showed a number of results. First, the quantitative phenotypes caused by the 50 mutations fell into clusters and subclusters. Significantly, almost all replicates of a given gene deletion grouped at least in the same subcluster, suggesting that differences among strains with different gene deletions were not the result of variability caused by experimental errors or artifacts. Second, the pathway and gene expression <u>output variables</u> (O and G) were sometimes affected by different genes than the <u>"cell-to-cell variability"</u> variables (η^2^(O), η^2^(G), η^2^(P)). For example, cluster II is comprised of all the entries with low pathway output (O) and low gene expression output (G). Within this cluster there are subclusters with high (IIa), low or unchanged η^2^(O) and η^2^(G) (IIb and IIc). Also note cluster III, which contains cases with high pathway (O) and gene expression (G) output and low cell-to-cell variability for pathway and gene expression. Within cluster III, subcluster IIIa is defined by low η^2^(P) and subcluster IIIb by high η^2^(P) at the low pheromone dose. Such genetic independence strongly suggests the existence of distinct mechanisms independently controlling the two types of quantitative phenotypes (output and variability) and disfavoring an interpretation in which variability is inextricably linked to output strength. Rather, output and variation in output emerged as independent axes of system behavior, subject to independent regulation, and independently affected by genetic changes (see Discussion).

Similarly, suppression of signaling variation at low and high system inputs required the action of different sets of genes. This is evidenced in the subclusters within cluster V. Cluster V groups datasets with high cell-to-cell variability in both system output and gene expression output. Subcluster Va contains datasets with high η^2^(P) at high pheromone stimulation and low η^2^(P) at low pheromone. In contrast those in subcluster Vb are high in η^2^(P) at high pheromone but high, low or unchanged in η^2^(P) at low pheromone. The opposite is true for subcluster Vc, whose members have high η^2^(P) at low pheromone but variable at high pheromone.

A second feature in the clustering results is that mutations in related genes cause similar patterns of change in their set of quantitative measurements. This was expected and reassuring. For example, deletions of duplicated paralogs of ribosomal protein genes are grouped in subclusters IIa and IIb (distinguished by their different variation phenotypes) and those for the two PRS MAPKs, *FUS3* and *KSS1*, are together in cluster IV. By analogy with dataset clustering studies based on gene expression data and other phenotypes, we expect that the gene deletions that shared a cluster or subcluster membership would share mechanistic functions in the control of quantitative phenotypes in the PRS.

Figure 6 presents an alternative representation of output variables and variability variables for the 50 strains. In it, we plotted O vs. P (estimated as O/G) at low dose (6a), η^2^(P) vs P at low dose (6b), and η^2^(P) vs P at high dose (6c). These results again show that different genes affect each of these quantities differently.

**Figure 6.**
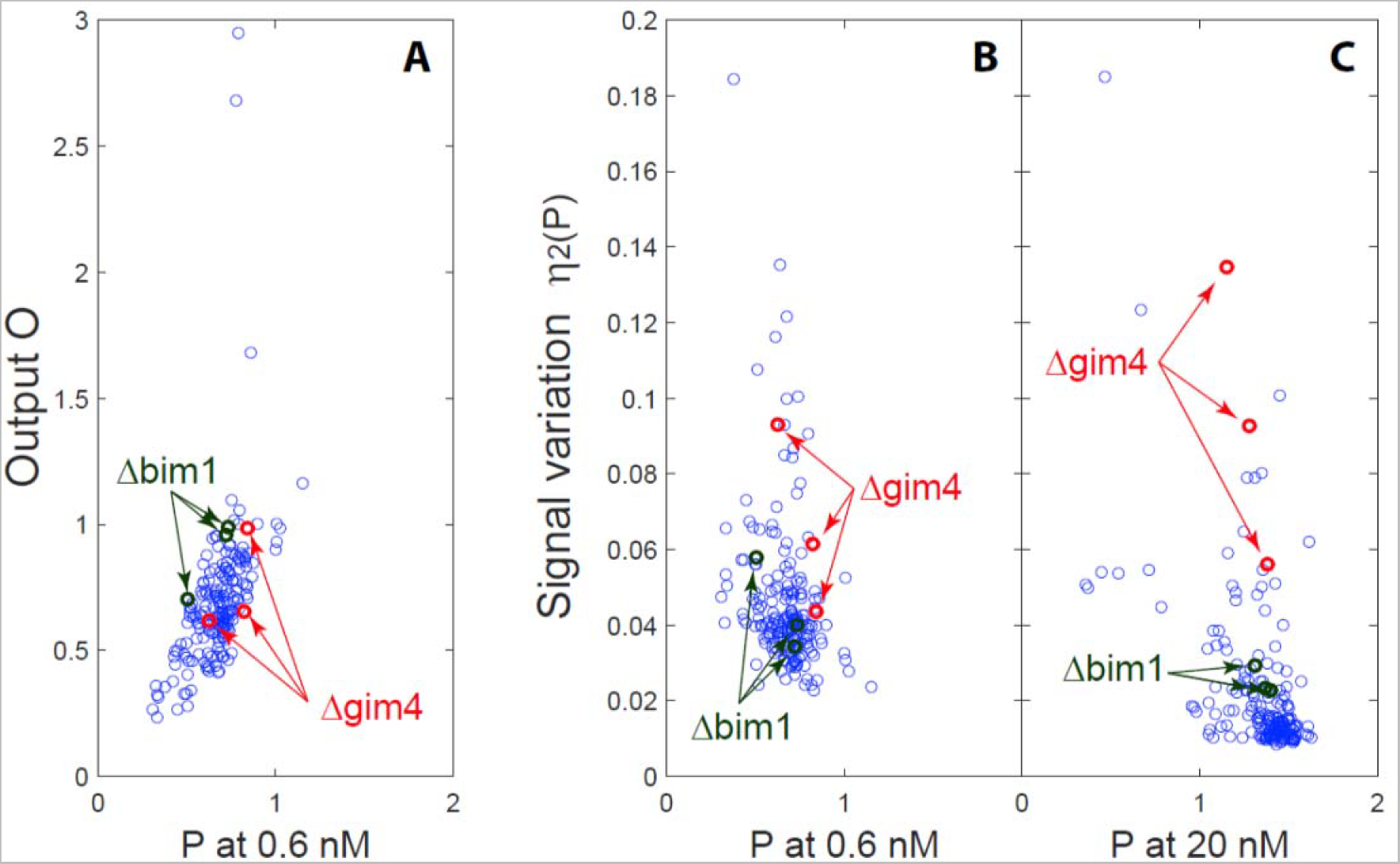
System output and signal variation vs. transmitted signal for signal and output variation strains. For the 50 mutants selected from the screen for further study, figure shows **(A)** System output O vs transmitted signal P (computed by dividing O by G. See Table 1) at low pheromone dose, showing good correlation/interchangeability of O and P, i.e. mutations that change the output do so by changing P, not G. **(B)** Signal variation η^2^(P) vs P at low dose and **(C)** η^2^(P) vs P at high dose. Arrows show Δbim1 and Δgim4, two mutations affecting microtubule function. At high dose both mutant strains show high η^2^(P) relative to the other selected strains. See Supplemental file Table_S3.xlsx for a list of all selected strains and their corresponding output and variation values.

### Two gene deletions with higher signaling variation affect microtubule function

Due to our interest in a possible model, in which the action of cytoplasmic microtubules decreased variation in transmitted signal by maintaining the cell nucleus at a constant position within a gradient of signal (discussed in Pesce et al. 2), we selected two genes affecting cytoplasmic microtubule function, BIM1 and GIM4 for further study. One gene, BIM1 affects attachment and function of the microtubule plus end to the signaling site. The other, GIM4, encodes a prefoldin subunit that affects microtubule function. We first verified the effects of these mutations by remaking them in a clean genetic background. To do so, we constructed a new reference strain, GPY4000 (see SI) that carried a pheromone responsive *P_PRM1_-mCherry* reporter and a second constitutive reporter, *P_BMH2_-YFP* reporter, and remade the deletion mutations in this clean background and again measured the variables defining quantitative system performance. Figure 7 shows average values of measurements by flow cytometry of η^2^(P) vs. O, i.e. of signaling variation vs. cumulative PRS output, of populations from the remade *Δbim1* and *Δgim4* single deletion strains, and a *Δbim1 Δgim4* double mutant. In these deletion strains, expression of the constitutive P_ACT1_ reporter was not affected (not shown). However, in these strains, signalin variation, η^2^(P), was increased similarly in both deletion strains, across all pheromone doses. Figure 7 shows this result, with η^2^(P) and system output O measured at different pheromone doses. This result is in contrast to our results in the original flow cytometric screen, in which *Δgim4* only increased η^2^(P) at low doses. In *Δbim1 Δgim4* cells, the increase in η^2^(P) was more than twice as large as the measured effect of the two individual deletions (Figure 7). This is thus an epistatic interaction (as defined by Fisher (1918)) suggesting that two gene products might have independent effects on signaling variation.

**Figure 7.**
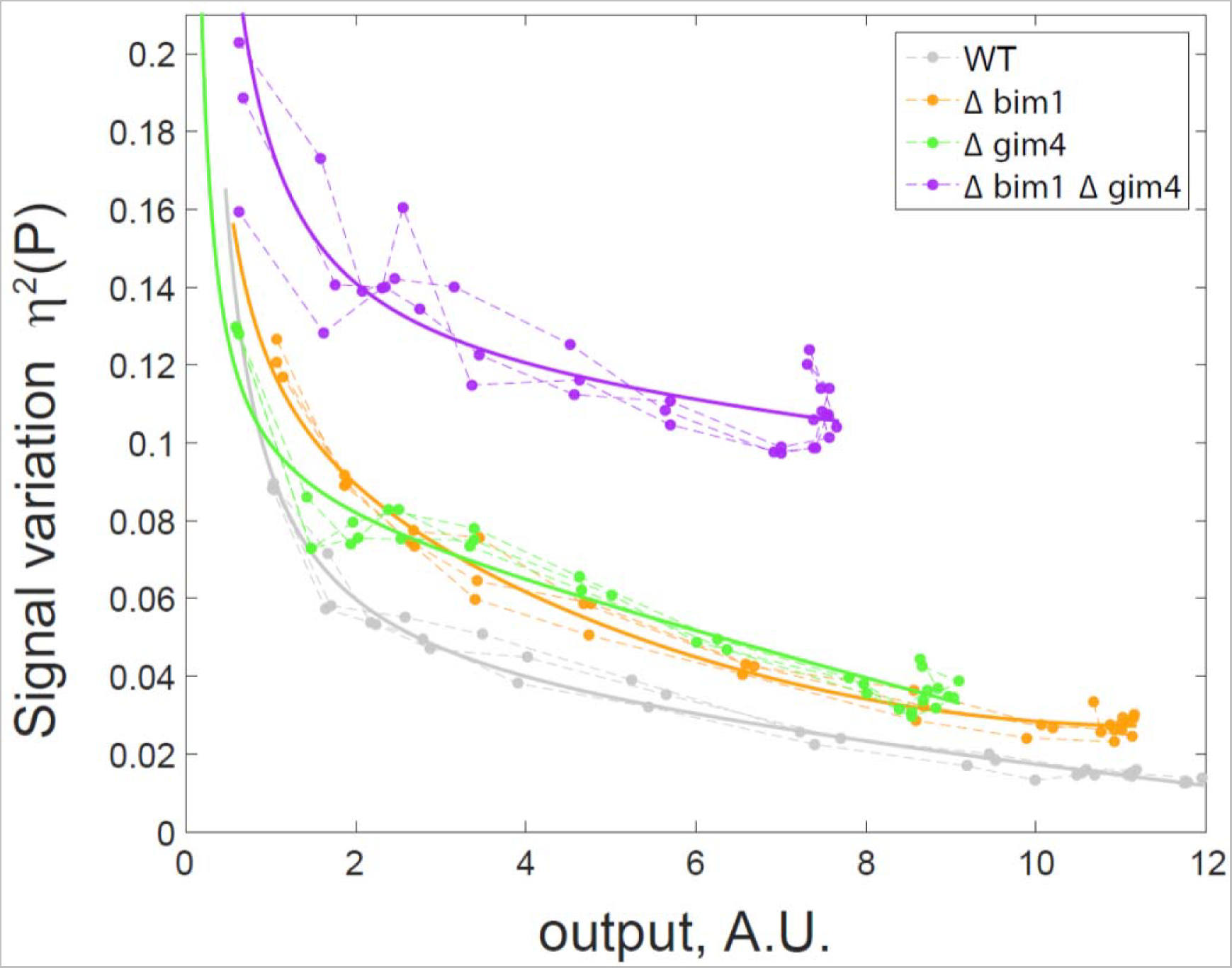
Deletions of *BIM1* and *GIM4* in clean genetic background increase signal variation at all outputs. Figure shows dose-dependent changes in pathway variation and cumulative transmitted signal in carefully reconstructed single- and double-deletion mutants. Plots show η^2^(P) as a function of O. Data were collected from dose response flow cytometry measurements of reference (GPY4000), *Δbim1* (GPY4001), *Δgim4* (GPY4031), and *Δbim1 Δgim4* (GPY4036) (bearing the *P_PRM1_-mCherry* and *P_BMH2_-YFP* reporters) stimulated for 3h with the indicated pheromone doses. Dashed lines connect measurements at different doses made on the same day. Solid curves drawn to show the best fit to a rational polynomial model for measurements from each strain. These models go to infinity at zero output, where we expect very large relative variation.

## Discussion

### Genes that suppress cell-to-cell variability in system output

We carried out a high-throughput screen for yeast genes whose products altered quantitative aspects of signaling and response in the yeast (PRS). Our approach required construction of thousands of mutant yeast strains, each of which carried multiple reporter genes and other alleles. This screen yielded mutants that affected system output, total transmitted signal, and cell-to-cell variation in those quantities. The fact that we found mutants that affected variation in transmitted signal is in contrast to some previous work, which found that deletion of genes affecting histone deactylation and ubiquitylation affected expression noise (Weinberger et al. 2012) affected expression noise from different promoters differently.

### High quality whole-genome variation deletion collection

We created a MAT α strain, SGA88, closely genetically related to strains in the haploid deletion collection (Chu and Davis, 2007) that carried reporters needed to quantify cell-to-cell variation in signaling and response, all linked to selectable markers. We mated SGA88 to a fresh instance of the original collection (a gift of Amy Tsong and Charlie Boone) to create a collection of more than 4,100 diploid strains (Pesce Heterozygote Deletion Diploid Variation collection (PHDDVC)). To use this PHDDVC collection, we sporulated each diploid strain on medium that selected for cells of the MATa mating type, for the deletion, and for the five additional genetic elements needed to carry out the assay. We then grew multiple individual cultures of clonal haploid cells from different small colonies arising from each meiosis. By passing the haploid deletions in the starting deletion collection through diploids and then sporulating those into a fresh haploid deletion collection containing our reporters, we ensured that consistently observed phenotypes were due to the deletions and not from unlinked mutations, including faster-growing mutations pre-existing in wells in the Chu and Davis collection (Hittinger and Carroll, 2007) or arising during meiosis. Moreover, by assaying cultures freshly derived from new meioses, we ensured the phenotypes derived from euploid cells that did not carry second site suppressor mutations, by avoiding contamination of slow-growing gene deletion mutant strains by faster growing aneuploid or second site suppressor variants (Hughes et al. 2000) and same-sex diploid variants (Giaever and Nislow, 2014) which are known to be present in wells of the Chu and Davis collection. This use of multiple freshly generated clonal cultures derived from independent meioses to diminish possible effects of unintended genetic heterogeneity is novel, and we hope relevant for subsequent quantitative studies of cell-to-cell variation and other subtle phenotypes measured in populations of different deletion mutants.

### High precision high-throughput screen

We then devised high throughxput means to search for mutants that affected cell-to-cell variation. These methods built on our previous microscopic cytometry work (Bush and Colman-Lerner; Colman-Lerner et al., 2005; Gordon et al., 2007), and on flow cytometry work by Newman and Weissman (Newman et al., 2006). By using flow cytometry, we compensated for differences in exponential growth rates of cells with different mutations, and accurately measured single-cell fluorescence from cultures that spanned a 10-fold range of cell concentrations. They included a number of steps to lower measurement errors sufficiently to unmask the effects of mutations on different components of variation. These included the use of multiple reporters with different colored readouts, ensuring that the cells were in exponential phase, paying careful attention to pheromone concentrations and to consistent conditions for inducing the PRS, collecting data from numerous independent cultures of control cells located randomly in different wells on every plate, and using cycloheximide to fix the cells and arrest protein translation, thus allowing time for all translated fluorophores to mature (Gordon et al., 2007) and also enabling us to “freeze” the assays at a precise time after system induction. By these means, we assayed the effects of more than 1100 deletions of nonessential genes, including strains bearing deletions in all 145 non-essential yeast protein kinases and phosphatase coding genes, and in 900 randomly selected genes. From these, we identified 50 genes that changed PRS output O, increased cell-to-cell variation in PRS output, η^2^(O), or increased cell-to-cell variation in signal transmission, η^2^(P). We hope that these methodological tricks might find use in other settings, for example in reducing measurement error and allowing dissection of sources of variation in studies of signaling and gene expression in mammalian cells directed toward genetic understanding, or toward identification of drugs (eg. drugs affecting gene expression from integrated proviruses such as HIV-1 (Dar et al., 2014)).

### Axes and epistasis groups

We clustered the 50 mutants under study into functional classes. To do so, we used a distance measure of each strain’s quantitative phenotypes including total PRS output, O, gene expression capacity, G, and variation in these numbers. This analysis showed that different sets of genes independently affected quantitative measures of transmitted signal, system output, and global gene expression capacity, and variability in these quantities. Further analysis showed that the effects of some mutants were specific to absolute level of these quantities, or to variation in them, establishing them as “axes” of quantitative system behavior whose values may depend on different cellular mechanisms. In their complexity, the results of this first order algorithmic classification were evocative of early-phase results of other mutant screens, for example the hunts for genes specifically required for anterior-posterior and dorsal ventral development in *Drosophila melanogaster* larvae (see Nüsslein-Volhard, 1979) and the assignment of DNA repair genes in yeast to different “epistasis groups”. In those cases, continued analysis led to simplification, and eventually to elaboration of the molecular mechanisms that determine the anterior-posterior and dorsal ventral axes in *Drosophila*, (Nüsslein-Volhard, 1987) and genetic and biochemical demonstration of independent mechanisms of DNA repair (Friedberg et al. 1995). In a companion paper, we discuss how our identification of genes that specifically affect η^2^(O) and η^2^(P) in the PRS adds to the list of other genes and loci recently identified in yeast, *Arabidopsis thaliana*, maize, and mice that reduce (canalize) the standard deviation of quantitative traits, rather than their mean values.

### Functional inference from whole gene deletion mutants and its limitations

Our work showed that two genes involved in microtubule function: *GIM4* and *PAC10*/*GIM2*, which encode members of the prefoldin complex, and *BIM1*, needed for cyto-plasmic microtubule dynamics, reduced cell-to-cell variation in signal transmission, η^2^(P). The fact that our genetic screen identified important players in this pathway, which has been so thoroughly studied, highlights both the sensitivity of the methodology we implemented, and the novelty of the quantitative phenotypes under study. It also underscores both powers and limitations of the genetic analysis based on whole gene deletion mutants: screening of deletion collections can uncover unexpected genes as key players in processes, but cannot always shed much light on their action. However, previous studies of cytoplasmic microtubules offer tantalizing clues to how these genes might act to reduce variation. During the pheromone response, a microtubule bridge connects the signaling site on the cell membrane with the cell nucleus, where genes are induced. Bim1 and Kar3/Cik1 attach microtubule plus ends to the signaling site and, by promoting polymerization and depolymerization, alternately push the nucleus away from the signaling site and pull the nucleus towards it. This alternation ensures that the nucleus localizes to a defined cellular location at the base of the mating protrusion (Maddox et al., 2000; Maddox et al., 2003). We wondered whether microtubule function might reduce variation in transmitted signal by mechanically positioning the nucleus within a gradient of signal originating at the signaling site. In a companion paper, we describe experiments to test this idea, and to determine the microtubule dependent process(es) necessary for reduction of signaling variation.

## Methods

General methods for cultivation of yeast strains and plasmid constructions are detailed in Supplemental Information.

### Analysis of cell-to-cell variation

We performed the analysis as in Colman-Lerner et al. (2005). Briefly, we considered the system output for any given cell O_i_, determined by the abundance of a fluorescent protein inducible by the pheromone response system, to be the product of i) the average pathway subsystem output per unit time, Pi (which varies with input pheromone dose), ii) the expression subsystem output Ei, and iii) the duration of stimulation ΔT (Colman-Lerner et al., 2005), as follows:

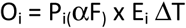

We considered P_i_ and E_i_ to be the sum of the capacity of the subsystem in each cell (Li and Gi) plus stochastic fluctuations in the operation of each subsystem during the course of an experiment (λ_i_ and γ_i_). Thus,

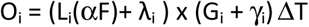

We defined the cell-to-cell variation in system output as the normalized variance of O_i_, η^2^(O), decomposable into the sum of individual sources a correlation term (Colman-Lerner et al., 2005), as follows,

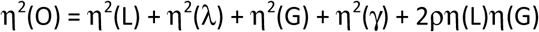

In the WT, in the deletion strains, and mutants used in the manuscript, we measured output and cell-to-cell variability of each reporter; η^2^(γ), gene expression noise ("intrinsic noise" (Elowitz et al., 2002)); as well as η^2^(P) (η^2^(L) + n^2^(λ)), cell-to-cell variability of the pheromone response system. We measured η^2^(γ) as the variance in the difference of the normalized abundance of the two fluorescent proteins driven by identical *P_PRM1_* promoters. We estimated cumulative signal transmitted P in individual cells as the normalized signal from the pheromone inducible *P_PRM1_* reporter (O) divided by the one the signal from the constitutive control promoter, *(P_ACT1_* or *P_BMH2_*, depending on the strain) (O/G). We estimated η^2^(P) as the variance in the difference between the normalized abundances of two fluorescent proteins, one driven by the *P_PRM1_* promoter and the other by the constitutive, pheromone-independent promoter (*P_ACT1_* or *P_BMH2_*, depending on the strain) (*σ*^2^(*mRFPj_i_*/< *mRFP* >−*YFP_i_*/<*YFP* >)).This variance is actually equal to η^2^(P)+ η^2^(γ), but η^2^(γ) was low enough in the WT and the 42 mutants in which we measured it (see SI) to assume that it may be neglected (Colman-Lerner et al., 2005).

### Construction of Heterozygous Diploid Deletion Collection and its use to generate sets of haploid deletion strains for screening

We constructed a MAT α strain, SGA88, which carried two pheromone inducible reporter genes, one constitutive reporter gene, a *bar1*-mutation which blocked a protease that removed pheromone from the extracellular medium, and a *cdc28*-*as2* mutation which allowed us to block the inhibition of the pheromone response by the cell cycle machinery by adding to the cells a chemical inhibitor of the mutant protein kinase. In SGA88 all of these genetic elements and the MATa marker were linked to individually selectable recessive (nutritional auxotrophy) or dominant (antibiotic resistance) markers. We mated SGA88 to a fresh instance of the original ("1.0") haploid deletion collection (Chu and Davis, 2007, a gift of Amy Chu) to create the Pesce Heterozygous Deletion Diploid Variation collection (PHDDV collection), comprised of more than 4,100 diploid strains. In these diploid strains, three dominant resistant markers: hygB^R^, G418^R^, Nat^R^, and two recessive markers His3 and Leu2, allowed selection of genetic elements, while two dominant sensitivity markers: Canavanine^s^ (due to the *CAN1* allele) and Thialysine^S^ (due to the *LYP1* allele) allowed selection against unsporulated diploids. We then sporulated different members of the PHDDV collection on appropriate selective medium to generate haploids that bore the deletion and the other genetic markers needed for the screen. We picked these as individual small colonies on selective plates and assayed individual cultures grown from these colonies.

To screen for mutants that affected cell-to-cell variation in pathway output, we grew cells in log phase (<3-10^6^ cells/ ml) for at least 14 hours. This step is in contrast to the standard practice of diluting carbon-exhausted cultures 4-6 hours prior to measuring them. By relying on exponential phase cultures we minimized undesired variability in PRS output arising from strain-to-strain and day-to-day differences in time to enter the exponential growth phase. We exposed our cultures for 3 h to two different pheromone concentrations (0.6 nM or 20 nM) and 10 μM *cdc28*-*as2* inhibitor 1-NM-PP1. We then added 50 μg/ml cycloheximide to inhibit protein synthesis and allowed for existing translated fluorescent protein molecules to mature (Colman-Lerner et al. 2005, Gordon et al. 2007). To aid the mutant screen and follow-up experiments, we measured the maturation times of mRFP (strain collection) and mCherry (follow up experiments) after blocking protein synthesis with cyclohexamide as in Gordon et al (Gordon et al., 2007) (not shown). Measured 1/2 time to maturation was 120 mins (RFP) and 45 mins (mCherry).

We measured fluorescent signal from the *P_PRM1_*-*mRFP* and *P_ACT1_*-*YFP* reporters by cytometry (BD LSRII with HTS auto-sampling attachment) and calculated or estimated parameters of interest, such as system output O_i_ and cell-to-cell variation in signal transmission, η^2^(P), as described above. We then verified (by cytometry) altered behaviors in three additional clonal isolates from the same mating, as described above. We confirmed by PCR in a random strain from the set of four for the presence of the expected deletion and the absence of the wild type coding sequence. We checked this strain by image cytometric fluorescent microscopy at the two different doses to confirm lack of aggregation and to measure *P_PRM1_*-*CFP* signal. Measurement of CFP signal allowed us to determine if the mutants affected η^2^(γ). As described, η^2^(γ) was a small contributor to cell-to-cell differences in gene expression and no mutant affected it.

## Author contributions

GP designed and constructed the strain collection, performed the screen, analyzed the results and characterized the selected mutants. DR assisted in this work. GP and SZ designed and performed the microtubule perturbation experiments. WP analyzed the numerical data in Figure 6D. RCY contributed to experimental design, data analysis and earlier versions of the manuscript. RB, GP and AC-L directed and guided the work and its interpretation. GP, AC-L and RB wrote the paper and guarantee the integrity of its results.

## Acknowledgments

We are grateful to Amy Tong, Charlie Boone, Guri Giaever and Corey Nislow for strains, plasmids, advice, and previously unpublished information for constructing our strain collection, Steve Andrews, Alexander Mendenhall, and Alan Bush for valuable discussions throughout. Work was supported by R01 GM097479 to RB. Earlier work received support from grants R01 GM086615 to RB and RCY and from P50 HG002370 to RB.

---

**Figure S1.**
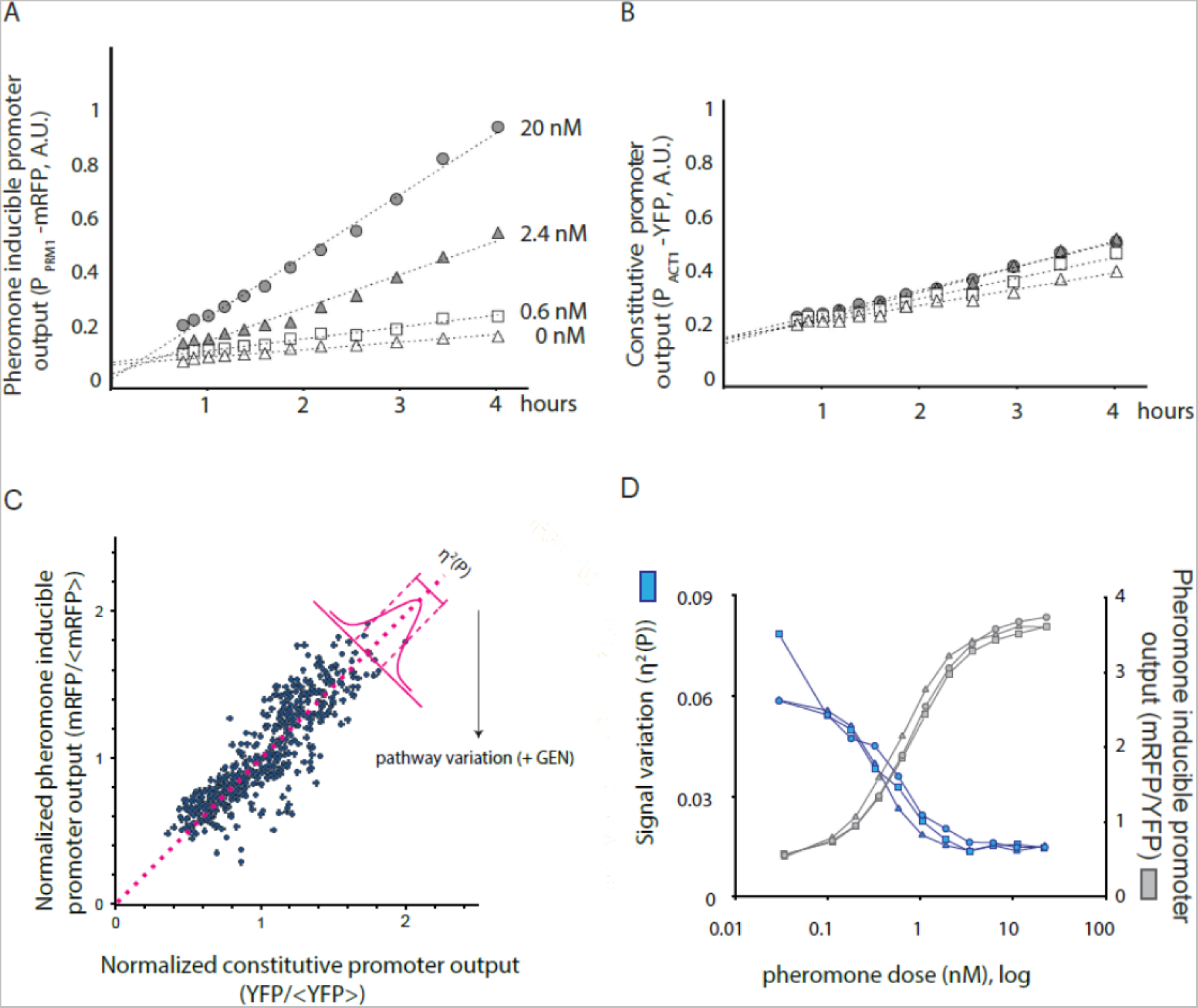
Time dependent output and Dose-Response of the reporter genes used in the screen. We stimulated SGA85 cells with the indicated concentrations of pheromone and measured the accumulated fluorescent protein by flow cytometry as detailed in Methods. **A, B.** Average fluorescence output of the pheromone-inducible *P_PRM1_-mCherry* reporter (A) and the constitutive *P_ACT1_-YFP* reporter **(B)**, in A.U., measured at four different doses over time. **C.** Estimating cell-to-cell variation in the activity of the pathway (η^2^(P)). Panel shows a plot of the pheromone-induced reporter *(P_PRM1_-mRFP)* output and a constitutive reporter (*P_ACT1_-YFP*) output in 500 cells of the reference strain (SGA85) stimulated with 20 nM pheromone for 3 h. For each cell, the amount of pathway variation (η^2^(P) + n^2^(γ)) (a quantity very close to η^2^(P), see SI) is the distance of each each cell from the identity line, drawn in pink. **D.** Dose-dependence of pathway variation and output. Plot shows pathway variation η^2^(P)+η^2^(γ) (blue), and output (grey) in SGA85 cells, as a function of pheromone dose after 180 min.

**Figure S2.**
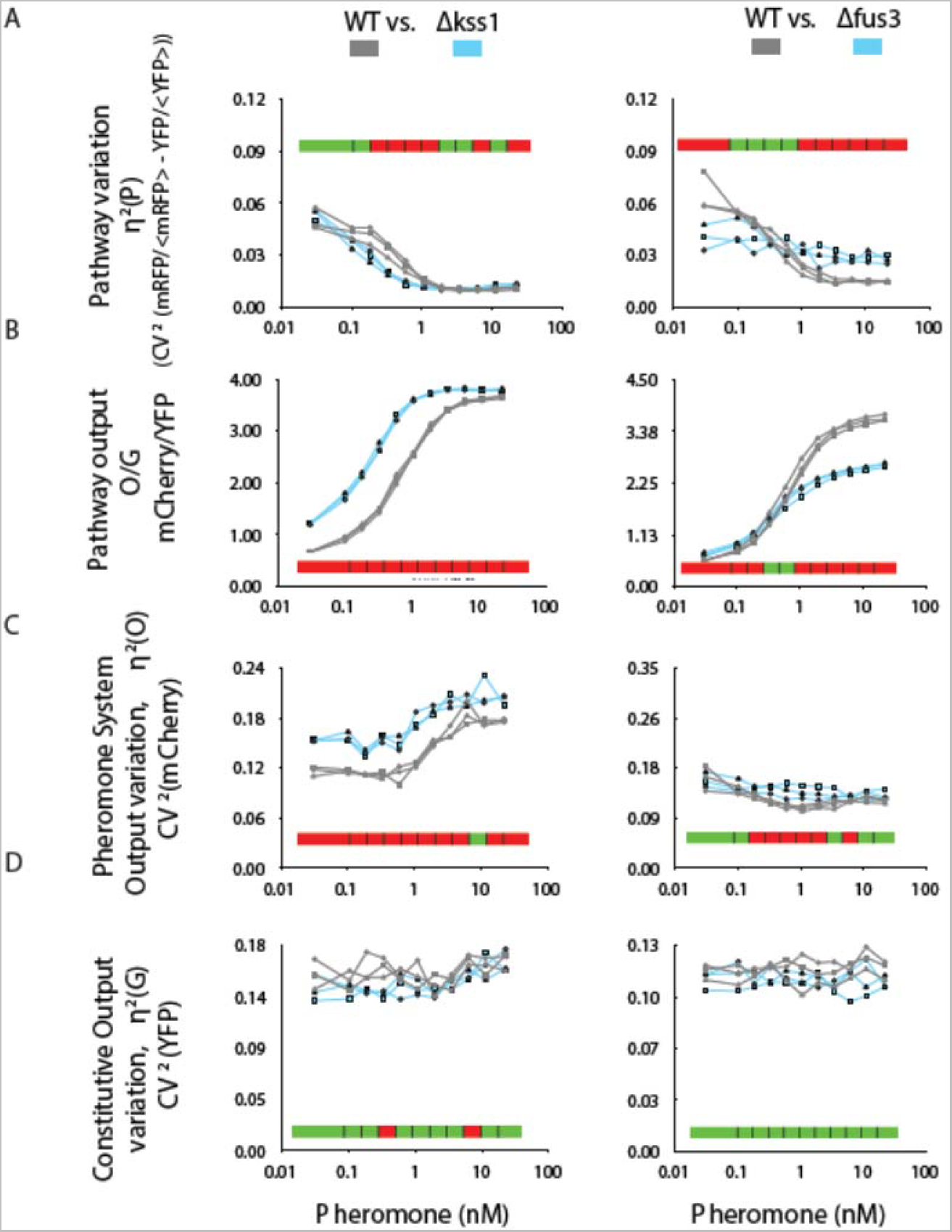
Genes encoding the MAPKs *KSS1* and *FUS3* affect pathway variation. Flow cytometry measurements of reference (here called “WT”, GPY4000, grey), *Δkss1* (GPY4038, left panels) and *Δfus3* (GPY4004, right panels) strains (bearing *P_PRM1_-mCherry* and *P_BMH2_-YFP* reporters), stimulated for 3 h with the indicated pheromone doses. Plots show values of different system measurements (see Table 1 and SI) vs. pheromone dose for three biological replicates of each strain. For each plot, red color along the horizontal bar indicates 0.95 confidence statistical significance of differences at the corresponding dose, as assessed by T test. Green color indicates no significance. **A.** Variation in transmitted signal (estimated by η^2^(P)+η^2^(γ)). **B.** Estimated P, that is, O/G **C.** Cell-to-cell variation in PRS system output, η^2^(O), measured as output of P_PRM1_-mCherry reporter. **D.** Cell-to-cell variation in gene expression capacity, η^2^(G), estimated by variation in output of the constitutive P_ACT1_ reporter.

## 1 Yeast strains

### 1.1 General methods

We performed DNA manipulation including PCR and subcloning as described (Ausubel et al., 1987-2016). We cultured and manipulated yeast as described (Ausubel et al., 1987-2006; Guthrie and Fink, 1991). Unless otherwise noted, we grew cells in synthetic dextrose complete (SDC) media consisting of Brent Supplemental Media (MP Biomedicals, Solon, OH), yeast nitrogen base without amino acids and ammonium sulfate (BD, Franklin Lakes, NJ), and dextrose (Sigma-Aldrich, St Louis, MO).

### 1.2 Strains

We describe constructions of these strains and plasmids below. We also list We list strains used in this study in Table S4.

#### 1.2.1 Construction of SGA88, the MATα partner to be mated with the deletion collection

We generated a library of haploid **MATa** gene-deletion mutants containing the modifications listed in Figure S1b. To do this we first constructed **MATα** strain SGA88 containing the genetic elements needed for our screen (P_PRM1_-RFP, P_PRM1_-CFP, P_ACT1_-YFP, *cdc28-as2* and *Δbar1*), each marked with a different selectable marker. SGA88 also had a “cassette” *(Δcan1::P_MFA1_-LEU2*) carrying a **MATa**-specific marker (*P_MFA1_-LEU2*) that allowed positive selection of descendant haploid **MATa** cells and two recessive drug resistant markers (*Δcan1* and *Δlyp1*) that allowed negative selection of diploids and parental **MATa** cells (see more on the selection process below).

We constructed SGA88 in two separate, parallel tracks. **Track 1** resulted in a **MATa** strain. **Track 2** began with and resulted in a **MATα** strain. We introduced a subset of the desired modifications into strains from these tracks and then combined them into a single haploid strain isolated after mating and sporulation.

##### Track 1. Construction of MATa parent of SGA88

###### Starting strain

The starting strain for Track 1 was **MATα** strain y3656 (a generous gift from Amy Tong and Charlie Boone) carrying a *Δcan1::P_MFA1_-HIS3–P_MFalpha1_-LEU2* cassette in the BY4742 (MATα) background. BY4742 is *his3Δ1 leu2Δ0 ura3Δ0 lys2Δ0*.

###### Replacement of the BAR1 ORF with a CFP pheromone-responsive reporter and a selectable marker

We constructed plasmid pBUPC by inserting a 200 bp cassette in the HindIII site of pRS406. This cassette was composed of 100 bp of BAR1 locus sequence immediately upstream of the start ATG codon (P_BAR1_), 100 bp of sequence immediately downstream of the stop codon (T_BAR1_) and a linker with an Ase I site in between them. The relevant segment of pBUPC had the following orientation:

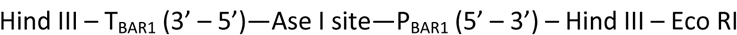

We then inserted a P_PRM1_-CFP-T_ADH1_ pheromone transcriptional reporter cassette in the marked EcoRI site of pBUPC, yielding plasmid pBUPC-PRM1-YFP. When cut with Ase I pBUPC-PRM1-YFP yielded the following linear DNA molecule:

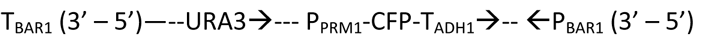

(arrows denote direction of transcription initiating at the noted promoters)

We transformed y3656 with AseI-linearized pBUPC to obtain strain SGA10. We confirmed proper integration at the *BAR1* locus by PCR and tested that the strain was phenotypically *bar1* and had a pheromone-inducible CFP reporter.

###### Replacement of the PRM1 ORF with a YFP pheromone-responsive reporter and a selectable marker

We introduced a second, different colored, pheromone-inducible fluorescent protein reporter in SGA10. To do this we replaced the PRM1 ORF with the following DNA piece obtained by amplifying the YFP-NAT resistance cassette from a plasmid from the *e collection* by PCR (Goldstein and McCusker, 1999; Longtine et al., 1998).

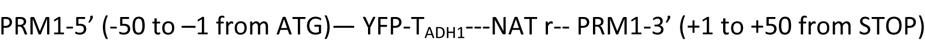
We transformed this PCR product into SGA10, selected for NAT resistance and confirmed integration by PCR. We also confirmed that the resulting strain had now both CFP and YFP pheromone inducible reporters. We named this strain SGA30.

###### Change of mating type to MATa

Our initial plan had been to introduce the reporters into a MATα strain that would be mated to the deletion collection. We realized that we could check that the reporters in such a strain were inducible by mating pheromone using **a** factor, but, because a factor is not very soluble and hard to work with, to calibrate the performance of the reporters for any screen, we would have to verify their performance in MATa. We therefore decided to calibrate the reporters carefully in such a MATa background before moving further. We therefore changed MATα reporter strain to a MATa. To do so, we mated the MATα SGA30 strain with a MATa strain from the BY4741 (MATa *his3Δ1 leu2Δ0 ura3Δ0 met15Δ0*)—derived deletion collection (we used the *Δace2::Neo r* strain). We did not name nor store the resulting diploid. We sporulated the it and by plating in the appropriate multi-selection plates, obtained a segregant with the following phenotype:

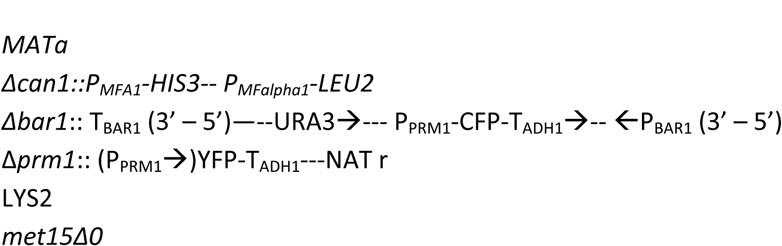

We named the resulting strain SGA31.

###### Switch of Abar1 marker from URA3 to HIS3

When making SGA31, we discovered that haploid colonies from the sporulated diploid growing in uracil/ histidine drop-out plates with NAT contained both uracil auxotrophic and prototrophic segregants. We hypothesized, and then demonstrated, that *ura3* auxotrophic segregants were able to scavenge uracil derived from their *URA3* neighbors in the colony. Since our plan for generating a modified deletion collection depended on our ability to stringently select only strains that carried the *Δbar1::P_PRM1_-CFP* reporter, we decided to not rely on *URA3* to select strains. We therefore resolved to replace the *URA3* marker for this reporter with a *HIS3* marker. We knew that *HIS3* allowed clean selections: we never found *MATα* segregants not expressing *P_MFA1_-HIS3* mixed in with the *HIS3*-expressing *MATa* segregants.

To switch the marker in *Δbar1* we first needed to switch the *P_MFA1_-HIS3–P_MFalpha1_-LEU2* to a different cassette that did not rely on *HIS3* expression to select *MATa* cells. To this end we obtained strain y3996 from Amy Tong and Charlie Boone. y3996 carried the following haploid-selection locus:

###### Addition of Δcan1::P_MFA1_-LEU2 marker

This haploid selection locus, by contrast to y3645’s, did not allow for selection of *MATα* segregants, but rather only *MATa* segregants. However, since selection of *MATα* segregants was not necessary for our approach, we switched to this simpler selection cassette, freeing the *HIS3* locus to be used to mark *Δbar1*. We therefore mated SGA31 with strain y3996 and used appropriate selection to identity a segregant, SGA33, with the following elements:

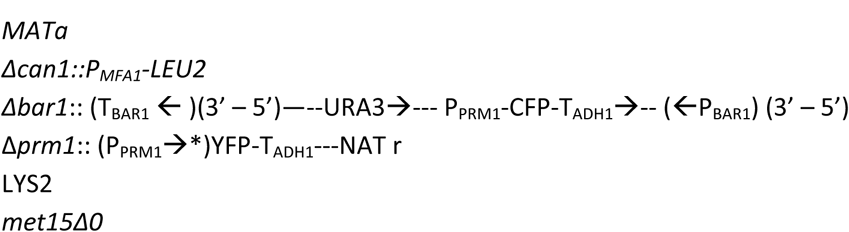

Here, the parenthesis around a series of symbols P_PRM1_, T_BAR1_ and P_BAR1_ indicate an endogenous element (promoter or terminator).

To replace the *URA3* marker in *Δbar1* with a *HIS3* marker we amplified the entire *HIS3* locus from pRS403 by PCR, using primers that yielded this PCR product along with 200 bp of pRS backbone sequences on each side. These same sequences flanked the *URA3* locus in the *URA3* marker inserted in *Δbar1*. We introduced this linear *HIS3* PCR product into SGA33 by transformation and selected for histidine prototrophy. All of the selected colonies had lost the *URA3* marker but had retained the pheromone-inducible *P_PRM1_-YFP* reporter. The resulting *Δbar1* locus was as shown in the schematics below.

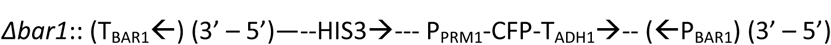

We named this strain SGA37.

###### Change in the DNA elements near the P_PRM1_-CFP reporter to correct for decreased total expression and greater expression variation caused by colliding polymerases

With a MATa strain in hand, we obtained high quality single cell level data of the expression of the CFP and YFP pheromone inducible promoters. We compared the results with those from a well-characterized control W303 background strain containing the same reporters, with the CFP reporter in a different locus and the YFP reporter in the same locus as in SGA37. We found that the CFP reporter, but not the YFP reporter, had reduced level of expression in SGA37. In addition, gene expression noise for the *PRM1* promoter was higher in SGA37 (Colman-Lerner et al. 2005). We hypothesized that these two differences might be due to transcription starting at the intact *BAR1* promoter clashing with convergent transcription starting at the *PRM1* promoter. We demonstrated that this was the case by deleting 300 bp of upstream *BAR1* sequences (the pheromone response elements in P_BAR1_ reach up to −273 from the ATG) and showing that the behavior of the _PPRM1_-CFP reporter changed to now match the behavior of the reporter in the W303 control strain.

To delete the *BAR1* promoter we amplified the *URA3* locus from pRS406 flanked with sequences from the upstream of *BAR1* and the terminator in the CFP reporter, as schematized in (1):

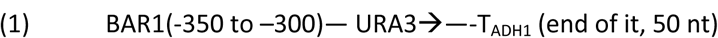

This PCR product was meant to recombine with the *Δbar1* locus in SGA37, schematized in (2):

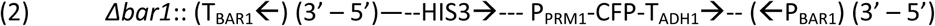

The final, modified *Δbar1* locus would have the structure shown in (3):

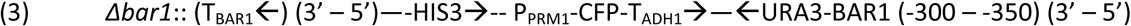

We transformed the PCR product depicted in (1) into SGA37 and confirmed proper recombination by PCR. The resulting strain was named SGA39. As stated above, the P_PRM1_-CFP-T_Adh1_ was restored to normal behavior by this procedure.

In a subsequent step we removed the URA3 marker by transforming SGA39 with a double stranded oligonucleotide with homology to the BAR1 (−300 −350) segment and the *URA3* gene terminator, and plating on 5-FOA containing plates. The resulting *Δbar1* locus kept the URA3 terminator as a safeguard against any runaway polymerase that might go through that segment. The locus is schematized below:

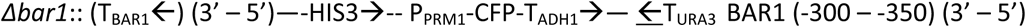

(the <u>←</u> denotes the direction of the URA3 terminator inserted, there is no known transcription starting in these sequences).

This locus is annotated as *Δbar1-ORF::PRM1pr-CFP–HIS3 Δbar1-promoter::URA3-terminator* in the strain table (Table S4).

The resulting strain was named SGA41.

SGA41 does not show the reduced expression and increased noise of the P_PRM1_-CFP reporter observed in parental strain SGA37. Both P_PRM1_ reporters in SGA41 showed results identical to the same P_PRM_i reporters in the reference W303 strain.

###### Introduction of a constitutive control reporter

At this point in the strain construction process, research in the W303 strain had uncovered that cell-to-cell variation in the single-cell levels of expression from any transcriptional fluorescent protein reporter was substantially increased by a large cell-to-cell variation in the capacity of cells to express genes into proteins (Colman-Lerner et al., 2005). In addition, the same work had developed an internal-reference method using an additional different-colored fluorescent protein transcriptional reporter driven by the promoter of a housekeeping gene. This method allowed the detection and quantification of η^2^(P), cell-to-cell differences in the strength of the signal received by the promoter. We therefore decided to alter our plans for the Track 1 strain to include a third fluorescent protein reporter, driven by the promoter for the housekeeping gene *ACT1*, which codes for actin.

At the same time, in parallel to these developments in our lab, the Boone lab had added a major improvement to the SGA pipeline by adding *Δlyp1* as a second selectable marker to eliminate unsporulated diploids in the haploid selection step (Tong et al., 2004). *LYP1* codes for a lysine transporter that is only required in lysine auxotrophs. The toxic lysine analog thialysine also enters the cells exclusively through the Lyp1 permease, therefore the *Δlyp1* mutation is a recessive thialysine resistance mutation. Addition of both canavanin and thialysine to the haploid selection plates provides a “double lock” mechanism to prevent the growth of unsporulated diploids.

We decided to combine our two new needs for a *Δlyp1* mutation and a constitutive fluorescent protein reporter by inserting the reporter replacing the *LYP1* ORF.

Similar to what we did above for *BAR1* and P_PRM1_-CFP, we constructed pLYP1a, containing a 200 bp cassette in the HindIII site of pRS406. This cassette was composed of 100 bp of *LYP1* locus sequence immediately upstream of the start ATG codon (P_LYP1_), 100 bp of sequence immediately downstream of the stop codon (T_LYP1_) and a linker with an Aflll site in between them. We then cloned a P_ACT1_-YFP cassette adjacent to the *LYP1* integration cassette, yielding pLYP1a-PACT1-YFP. The scheme below represents pLYP1a-PACT1-YFP linearized with Aflll

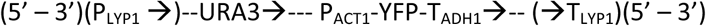

This locus is annotated as *Δlyp1::ACT1pr-YFP–URA3* in the strain table (Table S4).

Note that in contrast to the pBUPC construct used for *BAR1* and P_PRM1_-CFP, in the *LYP1* P_ACT1_-YFP construct the coding strands are the same for *LYP1*, *URA3* and *ACT1* sequences, preventing the “collision” of polymerases that affected the P_PRM1_-CFP reporter (see above).

We transformed Aflll-linearized pLYP1a-PACT1-YFP into SGA41 and selected for uracil prototrophy, yielding SGA43. We confirmed the correct insertion by PCR, by the acquisition of resistance to thialysine and the presence of a constitutively-expressed YFP.

##### Track 2. Construction of the MATa parent of SGA88

###### Starting strain

The starting strain for Track 2 was **MATα** strain BY4742. As described below, we introduced in this strain a *cdc28*-*as2* allele and a *P_PRM1_-mRFP* reporter. We later combined these two components of our final strain with components described above by mating and meiotic segregation.

###### Insertion of a selectable cdc28-as2 allele

We replaced the CDC28 gene with the analog-sensitive *cdc28-as2* allele using plasmid pCDC28-as2-406 (Colman-Lerner et al., 2005). Linearized pCDC28-as2-406 recombines with the CDC28 gene and integrates by a single recombination step.

The modified *CDC28* locus is schematized below:

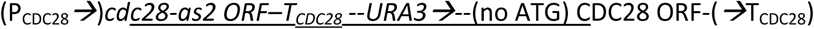
(plasmid sequences underlined. Note that the “(no ATG) *CDC28* ORF” doesn’t have a promoter upstream, besides not having a start codon).

We transformed linearized pCDC28-as2-406 into BY4742 and selected for uracil prototrophy. The resulting strain SGA60 was sensitive to the ATP analog 1-NM-PP1.

In our previous use of this construct we followed this transformation by selecting for “looped out” variants using 5-FOA selection, leaving an unmarked P_CDC28_ **→***cdc28-as2* locus. For the purpose of our SGA project we needed to have a selected marker next to *cdc28-as2*, a marker different than the ones used before (URA3 was occupied by the *Δlyp1* P_ACT1_-YFP element). We also needed to use a *cdc28-as2* element unable to revert to wild type *CDC28* by loop-out recombination.

We thus replaced the URA3 marker and the WT *CDC28* ORF and terminator homology at the 3′ end of the modified locus using a PCR approach with a hygromycin B resistance cassette schematized below:

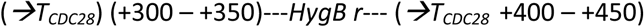

After a double recombination event with the PCR product, the structure of the *CDC28* locus would be as schematized below:

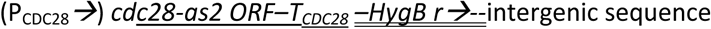

(PCR product sequences double underlined)

This locus is annotated as *Cdc28-F88A–HygB(MX4)* in the strain table (Table S4).

We transformed the *HygBr* PCR product into SGA60 and selected for hygromycin resistance to obtain strain SGA62. We confirmed proper insertion by PCR and by loss of uracil prototrophy. SGA62 was sensitive to 1-NM-PP1.

###### Insertion of a P_PRM1_-mRFP reporter

We constructed the *P_PRM1_-mRFP* reporter using a PCR approach as used above for the *P_PRM1_-YFP* reporter, replacing the *PRM1* ORF with the *mRFP* ORF followed by a selectable marker. We obtained an mRFP variant of the *Pringle collection* plasmids carrying the nourseothricin resistance (NAT r) selectable marker from the O’Shea lab (UCSF, (Huh et al., 2003)).

After transformation the modified *PRM1* locus has the structure schematized below:

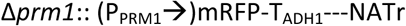

We transformed the mRFP-NATr PCR product targeting *PRM1* into SGA62 and selected for NAT resistance. We confirmed integration by PCR and by the presence of a pheromone inducible RFP. The resulting strain was named SGA64.

This locus is annotated as *Δprm1::PRM1pr-mCherry–G418(MX6)* in the strain table (Table S4).

###### Generation of SGA88 from Track 1 and 2 strains

We mated strains SGA43 and SGA64, yielding diploid DSGA1. We sporulated DSGA1, dispersed the spores and plated them in arginine and histidine dropout plates with canavanine, NAT and HygB. We replica-plated the resulting colonies in single-dropout plates for leucine, uracil, lysine and methionine.

We screened the uracil and methionine prototrophs and lysine and leucine auxotrophs (leucine auxotrophy identifies MATalpha cells) to find segregants with a pheromone inducible RFP reporter (since half would have it, the other half would have the YFP variant).

The resulting strain **SGA88** had the following genetic elements:

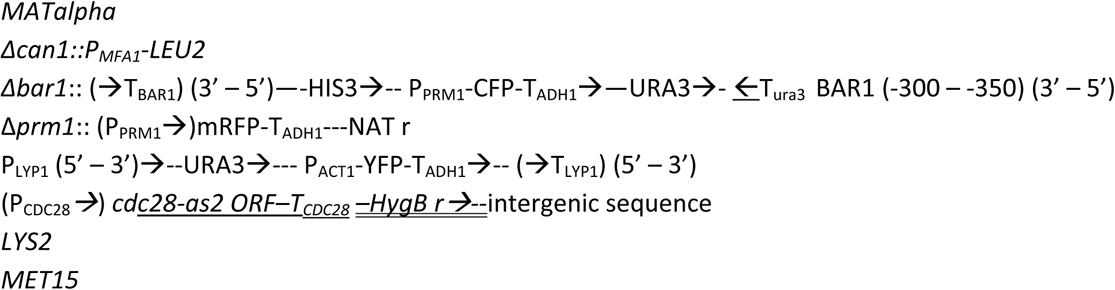

We tested for the presence of each of these elements in SGA88 by PCR, and also using phenotypic assays when available.

We used SGA88 as the MATα partner to mate with the non-essential haploid deletion collection.

As a control MATa strain, we also selected a sibling of SGA88 that grew without leucine (and thus was MATa) and that had an pheromone-inducible mRFP reporter. This strain was *met15Δ0* and was named **SGA85.** SGA85 became the control strain in the high throughput screen, seeded in multiple wells in each screening plate, and became the reference strain for follow up studies.

#### 1.2.2 Construction of a gene deletion library containing required reporters and selectable markers

We constructed a ˜4,000 strains gene deletion library in which each strain contained the genetic elements described in Figure S1B. We developed a modification of the procedure described by Charlie Boone and collaborators (Tong et al., 2001; Tong et al., 2004). See schematics in Figure S1A.

##### Mating of SGA88 with the deletion collection, and subsequent sporulation

Briefly, we first grew a lawn of the MATalpha partner SGA88 on rectangular YPD agar plates, pinned a freshly grown 384-format array of the yeast deletion collection on top of the lawn and incubated the plates for 24 h at 30° C. We next pinned the grown, partially diploid colonies onto diploid selection plates and incubated for 2 days at 30°C; this step eliminates the unmated haploids. We subsequently pinned the diploids onto sporulation plates and incubated for 5-8 days at 25°C. We monitored the progress of sporulation by counting the fraction of spore-containing asci in the colonies in the microscope. Once the number of asci was stable we stored the sporulated colonies at 4°C until the next step. The differences with the Tong and Bonne procedure are described below:

##### Diploid selection

We used antibiotic selection in G418 and HygB plates instead of relying on the auxotrophic markers *lys2Δ0* and *met15Δ0*.

##### Haploid selection

To isolate single haploid deletion strains carrying the above reporters, we sporulated the diploid colonies and manually streaked from these onto haploid selection plates instead of pinning them as a patch on the selection agar (see next section).

We used haploid selection medium that allowed us to positively select for a larger number of genetic elements than the process described by Tong and Boone (see next section). The negative selection for diploids in this medium was effected by canavanine (to kill all *Δcan1/CAN1* diploids) and thialysine (to kill all *Δlyp1/LYP1* diploids).

##### Selection of clonal haploids and assembly of the library for the flow cytometry-based screen

Because our screening method was aimed at detecting changes in standard deviation in single cell distributions we used a very stringent approach to collect the strains that made up our library. We decided against relying on culturing cells for the screening that had been selected in bulk from the colonies in the sporulation plate, which is still the standard approach in this type of high throughput methods of library construction (Tong and Boone, 2007).

Instead, we manually streaked each sporulated colony onto haploid selection media as described above.

Figure 2 in the main text shows the genetic markers haploid strains needed to have to grow on these plates.

To maximize the chances that all the cultures in our screen would be genetically homogeneous, we manually picked colonies from these plates. In several cases we found two or more populations of colonies on haploid selection plates distinguished by their size. We interpreted this size heterogeneity as a sign that the original gene deletion colony in the library carried a genetic change that suppressed a growth defect caused by the gene deletion (for example aneuploidy (Hughes et al., 2000), other second site suppressors (Hittinger and Carroll, 2007) and formation of same sex diploids (Giaever and Nislow, 2014). To minimize the presence of such mutations in our manually isolated clones, we avoided the larger colonies and picked only colonies from the smaller size subpopulation.

We inoculated each selected colony into liquid medium that contained the same selection agents as the haploid selection plates (Figure S1B). We used 500 μl of media in 1.1 ml capacity “deep well” polypropylene 96-format plates. In each plate we also included the following controls:

SGA85: 4-6 wells per plate

BY4741: 4 wells per plate.

We distributed the placement of the control cells on the plates used for the screen in random patterns.

We then grew the 96-well plates to carbon exhaustion, 2 days at 30°C. We stored half of the saturated cultures at 4 °C until the next step. We added 15% glycerol to the other half and preserved at −80 °C.

#### 1.2.3 Reference strains used for controls for genetic perturbations in followup studies

##### Base strain with constitutive reporter (GPY4000)

We used another strain to generate gene deletion variants for confirmation and follow up experiments (see below). To construct it we introduced a different-color reporter driven by the housekeeping gene *BMH2*. We first generated a plasmid carrying the reporter construct by modifying plasmid pTC-P_BMH2_-YFP-T_ADH1_-URA3 (Colman-Lerner et al., 2005): we removed its *URA3* marker and replaced it with a genomic PCR product of the *MET15* gene, generating plasmid pTC-P_BMH2_-YFP–T_ADH1_-MET15. We linearized pTC-P_BMH2_-YFP-T_ADH1_-MET15 with StuI, which cuts within the P_BMH2_ element, and transformed the linearized plasmid into GPY1804. We screened 8 methionine prototrophic (MET+) transformants using fluorescence microscopy to identify an integrant with a single copy of the P_BMH2_-YFP reporter. We named this strain **GPY4000** and annotated this locus in the strain table as *BMH2::BMH2pr-YFP–MET15* (Table S4).

##### Base strain derived from SGA85 (SGA101)

We used this strain to introduce alleles by loop in/ loop out replacement with *URA3*-marked plasmids and for the three-color FP live cell time courses (see below). Starting with the reference strain SGA85, we derived a *Δura3* strain by transforming SGA85 with a PCR product spanning the entire *LYP1* locus from promoter to terminator, amplified from BY4741 genomic DNA. We selected for loss of the *URA3* marker in 5-FOA plates and confirmed restoration of the *LYP1* ORF by verifying the loss of YFP fluorescence from P_*ACT1*_-YFP-T_*ADH1*_ and by diagnostic PCR. The resulting strain was a *URA3 LYP1* derivative of SGA85. We named it SGA101.

#### 1.2.4 Strains used for follow up studies

##### A) Gene deletion strains

We introduced all gene deletions in the GPY4000 strain by a PCR approach. We first obtained strains carrying the gene deletion of interest marked with G418 resistance from the haploid deletion collection. We then changed the G418 resistance to either NAT resistance or hygromycin B resistance by transforming the deletion strain with a PCR product of the resistance cassettes and selecting for the marker carried in the PCR product. Since all three drug resistance markers contain the same 300-400 bp promoter and terminator, those flanking sequences directed a double homologous recombination gene replacement. Proper recombination was confirmed by the loss of the original G418 resistance. Subsequently, we amplified the gene deletion locus by PCR, including 300-500 bp of 5′ and 3′ gene-specific flanking sequences in the amplicon. We transformed this PCR product into GPY4000, selected for the appropriate drug resistance and confirmed proper gene deletion by PCR. The resulting strains are listed in Table S2.

## 2 High throughput growth, induction of PRS, and flow cytometry screen

### 2.1 High throughput growth and pheromone treatment of 96-format arrayed yeast colonies

As explained above, we stored at 4°C saturated cultures of 96-format clones from the modified deletion collection described in Figure 2a. To screen, we followed the procedure shown in the schematic in Figure 2b and described below:

#### Growth to exponential phase

We used a slotted pinning tool to inoculate 5 μl of the saturated cultures stored at 4°C into 500 μl of SDC media in 1.1 ml-capacity polypropylene 96-well plates and grew these cultures to carbon exhaustion, 2 days at 30 °C. We then inoculated 5 μl of the freshly saturated cultures into 250 μl of SDC in 300 μl-capacity polycarbonate 96-well plates, grew them for 8-10 h at 30 °C, and measured and saved the 0D600 of all wells using a multiwell spectrophotometer. We used the measured OD information to calculate the dilution of inoculum needed to have most of the strains in exponential phase after 12-18 hours of growth. We then prepared several 300 μl-capacity polycarbonate plates as before and inoculated them using the 5 μl slotted pinning tool, at the calculated dilution, and 0.5, 1.5 and 3 times that amount.

After 15 h of growth, we measured 0D600 of all plates and used the OD data to choose the plate in which the largest number of strains 1) was in exponential phase and 2) had enough cells to be used in the next step.

Immediately prior to stimulation, we sonicated the culture flat-bottom plates by “floating” the plates in the sonication bath of a S-3000 MP Misonix sonicator (Misonix Inc), set at power 10, for 2 minutes in two periods of 1 minute each with a 30 sec rest in between periods.

We next inoculated 5 μl of the sonicated cells into 250 μl of pheromone media in 300 μl-capacity polycarbonate 96-well plates. We followed the conditions described previously (Colman-Lerner et al.,2005). Briefly, stimulation media contained pheromone in SDC media containing 20 μg/ml caseine (SIGMA), to block pheromone binding to the plastic walls, 5-10 μM 1-NM-PP1, to inhibit Cdc28-as2 and 0.15 X strength PBS to buffer pH and thus prevent caseine precipitation.

We incubated the cells in pheromone-containing medium for 3 h at 30 °C for the screen and for most experiments, except when indicated. The 30°C incubation was done in an air heated ˜30 cm rotation radius shaker. After the end of the pheromone incubation period, we added 100 μg/ml cycloheximide to stop reporter accumulation and allow for complete fluorophore maturation. Yeast cells in cycloheximide retain their shape and external appearance for more than 10 hours at 30 °C.

Finally, we sonicated the plates as described above and measured fluorescent protein expression by flow cytometry.

### 2.2 Primary screen, selection of mutants and follow-up studies

#### 2.2.1 Primary screen

We used the high-throughput growth, stimulation and flow cytometry process described above to screen two sets of gene deletion mutants from the library we generated.

**Set 1: Unbiased Genes**—This set consisted of 996 gene deletions randomly selected from the library. We made this set by picking clones from the library arrayed in 384 colonies format in order of appearance, starting in position A1, completing each row, and following with the next row. When a colony was missing we looked for a colony corresponding to the same gene deletion in the section for duplicates in the library, and added it if present. This selection was unbiased with respect to, among other things: gene location in the genome, gene ontology, name, ORF number and any phenotype of the deletion strain.

**Set 2: Kinases and Phosphatases**—To assemble this set we searched the Saccharomyces Genome Database (SGD) for genes annotated with “viable systematic deletion phenotype” and with the “function” GO terms “protein kinase activity” and “phosphoprotein phosphatase activity”. We retrieved 106 and 41 hits, respectively. Except for one gene in each set, all of these putative or confirmed protein kinases and phosphatases were represented in our modified gene deletion library. The total number of strains in this set was thus 145 (105 for protein kinases and 40 for phosphatases) Table S1 contains the list of all strains screened.

#### 2.2.2 Selection of mutants from primary screen

We selected gene deletion strains for follow up studies based on their pathway variation and pathway output values as described in the main text. We used as reference the values obtained from the included unmodified reference strains (SGA85, present in 4 wells in each plate) and the overall distribution of values for all the strains, both deletions and controls. Based on these two criteria we defined thresholds to select the mutants (Figure 4a, 4b, and 4c) so that the mutants and approximately 10% of the SGA85 controls had values outside the selected thresholds. We thus selected from each set.

**Set 1 (unbiased genes):** 102 of 996 gene deletion strains.

**Set 2 (kinases and phosphatases):** 38 of 145 gene deletion strains.

Given that we had set thresholds so that approximately 10% of the SGA85 strains would also be selected, we expected that approximately 10% of the selected genes would not show a reproducible difference from reference when re-assayed.

#### 2.2.3 Secondary screens

##### Repeated assays on the same independently isolated segregants

We included this step only for the gene deletions selected in **Set 1.**

We consolidated all 102 strains selected by their variation (see above) in 2 new culture plates with fresh media, grew them to saturation and assayed as described above for the primary screen.95 of the 102 re-tested deletion strains showed a changed above or below the thresholds and were selected for the next round of follow up.

##### Assays on three independently isolated segregants

We applied this next step to all **Set 1** candidates that showed reproducible results in the repetition above and to all **Set 2** candidates.

We assayed three new segregants from the same haploid selection plates in which the sporulated SGA88 x deletion strain diploid had been streaked. We picked three new colonies and inoculated them in fresh media in 96-well culture plates. We applied the same criteria as in the first pick, selecting colonies of representative size and avoiding colonies that were unusually large. For each deletion strain we grew four cultures in these new plates: three cultures corresponding to the newly isolated segregants and a fourth culture inoculated from the previous culture grown for the primary and first secondary screens. Each plate also included 4 to 6 wells with SGA85 cultures (reference strains).We found the following number of strains that passed this test (cell to cell variability in signal transmission η^2^(P), or total system output from the pheromone-inducible reporter were above or below the distribution of values of the reference strain cultures):

**Set 1**–37 of 95

**Set 2**–17 of 38

**Total**–50 of 133

We confirmed by PCR that all these strains carried the gene deletions attributed to them. The measured values and description of the 50 gene deletions that yielded reproducible results is shown in Table 1 and Table S2.

##### Microscope cytometry assays for gene expression noise and morphology

We performed a follow up assay using fluorescence microscopy. This assay allowed us the CFP fluorescence signal accurately, which we could not accomplish using flow cytometry setup. In our gene deletion library CFP is driven by P_PRM1_ integrated at the BAR1 locus; P_PRM1_ also drives mRFP in the *PRM1* locus. With this pair of reporters we could measure gene expression noise η^2^(γP) from the P_PRMi_ promoter (Colman-Lerner et al., 2005). In addition to providing gene expression noise values, the microscopy assays allowed us to verify that any increase in pathway variation was in fact due to differences in reporter-derived cytosolic fluorescence between isolated, live cells and not a secondary consequence of cell aggregation, unusual shape, autofluorescence specks or other interfering factors.

##### Protocol for microscopy assay

All strains were grown in test tubes with SDC in two phases. First, the strains were inoculated from petri dishes, streaked no more than 10 days earlier from a frozen stock, and cultured for 6-10 h at 30 °C. These pre-cultures were diluted into new test tubes with SDC, at different densities, calculated based on the known doubling time of each strain, and cultured for 15-20 h at 30 °C. Cultures with OD_600_ between 0. 2 and 1 were used for the microscope assay. We measured 45 strains with gene deletions, including the 44 listed below in Table 1, plus reference strains. Table S6 contains all data from the microscope cytometry screens.

We sonicated cells in eppendorf tubes and incubated them for 3 hours at 30 °C in media with 10 μM 1-NM-PP1 and 0, 0.6 nM or 20 nM α-factor. Cell densities in the assays were in the OD_60_0 0.02 −0.1 range. At end of the 3 h incubation cycoheximide was added to a final concentration of 100 μg/ml and samples were incubated at 30°C for 5-7 h. Cells images were obtained and analyzed as described previously (Colman-Lerner et al., 2005; Gordon et al., 2007).

**Table 1.** Strains from flow screens tested by microscope cytometry.

**SI Table 1. Strains from flow screens tested by microscope cytometry.**
| Unbiased screen |  |  |  | Kinase and phosphatase screen |  |  |  |
| --- | --- | --- | --- | --- | --- | --- | --- |
| 1 | *MSH1* | 12 | *RPL34A* | 1 | *BUB1* | 12 | *OCA1* |
| 2 | *ARG82* | 13 | *RPP2B* | 2 | *CLA4* | 13 | *PBS2* |
| 3 | *BIM1* | 14 | *SLA1* | 3 | *CKB2* | 14 | *PKH1* |
| 4 | *DEP1* | 15 | *SUM1* | 4 | *CTK1* | 15 | *PPG1* |
| 5 | *ERD1* | 16 | *SWI5* | 5 | *ELM1* | 16 | *PPZ1* |
| 6 | *FUS1* | 17 | *SXM1* | 6 | *FUS3* | 17 | *PRR1* |
| 7 | *KAR4* | 18 | *UBA4* | 7 | *GAL83* | 18 | *RCK1* |
| 8 | *MKC7* | 19 | *UME6* | 8 | *HOG1* | 19 | *SAP155* |
| 9 | *PAC10* | 20 | *VPS64* | 9 | *HSL1* | 20 | *SKY1* |
| 10 | *RPL12A* | 21 | *YHR189W* | 10 | *KIN3* | 21 | *SSK2* |
| 11 | *RPL19B* | 22 | *YKL137W* | 11 | *KSS1* | 22 | *YVH1* |

## 3. Pheromone dose-response experiments

For follow up studies on the gene deletions of interest and/or all other perturbation experiments, we adapted the high throughput protocol we used for the screen. The only modification was in how we cultured the cells. Instead of culturing them in deep-well 96-well plates, we used the test tube protocol described above for the fluorescence microscopy secondary screen. We did the pheromone stimulation step exactly as in the assays we used in the screen, in shallow-well 96-well plates, with the only difference being the addition of pheromone, which in this case was done using a pipette from a serial dilution of pheromone stocks instead of the slotted-pin tool we used for the screen. All dose response experiments were measured by flow cytometry as described above.

## 4. Flow cytometric measurements

We used a Becton-Dickinson LSRII flow cytometer equipped with a 100 mW 488 nm laser and a 150 mW 532 nm laser, as schematized in Figure S2-B. All filters and dichroic mirrors were from Chroma. We used a threshold value of forward scatter (FSC) from the 488 nm laser to trigger data collection. We calibrated threshold values of FSC to detect the smallest cells in an exponentially growing culture of wild type or mutant yeast cells. We measured YFP and mRFP (or mCherry) fluorescence from the light emitted during the 532 nm excitation, which was channeled, using mirrors, into an octagonal array of 8 photomultiplier tubes (PMTs), labeled A to H. Light entering the array is first split by a 735 nm long-pass dichroic mirror (735LP) and then split again by a 640 nm long-pass dichroic mirror (640LP). The light that goes through the 640LP dichroic is filtered through a 675 nm band-pass filter of 50 nm wavelength width (675/50) before hitting the B PMT. The lower wavelength light that reflects in the 640 LP dichroic is directed towards a 600LP dichroic, the reflected light is split by a 540LP dichroic. The light that passes the 540LP dichroic is filtered through a 550 nm band-pass of 10 nm width (550/10) before hitting the D PMT.

We took the signal from the B PMT as the fluorescence from mRFP (or mCherry). Cells expressing only YFP or CFP show the same signal in this channel as wild type cells. We took the signal from PMT D as the fluorescence from YFP. Cells expressing only mRFP or CFP show the same signal in this channel as wild type cells. We also took a signal for CFP, but this channel suffered from high background autofluorescence (largely from intracellular NAD(P)H UV-excited, cyan-emitting fluorescence) and was not useful for this project (we instead measured CFP expression by quantitative microscopy).

## 5. Derivation of formula for estimation of signal variation

Here we present the derivation of the formula for signal variation *η*^2^ (*P_y_*). In Colman-Lerner et al. 2005, we showed that the cell-to-cell variation observed in fluorescent reporter expression can be split into four contributions: (1) variation in signaling *η*^2^ (*P*), (2) variation in gene expression capacity *η* ^2^ (*G*), (3) stochastic fluctuations in gene expression *η*^2^ (*γ_y_*), and a correlation term 2*ρ*(*L*,*G*)*η*;(*L*)*η*(*G*), where*ρ*(*L*,*G*) is the correlation between mean signaling capacity and mean gene expression capacity, computed over the population. In a case where we used an inducible yellow reporter and a constitutive cyan reporter, labeling the corresponding quantities with subscripts *y* and *c*, we would thus have

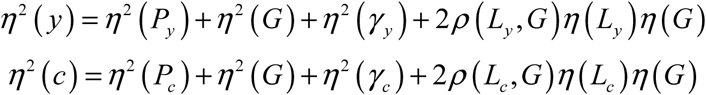

The measured correlation between these two reporters can also be split into contributions from the two subsystems:

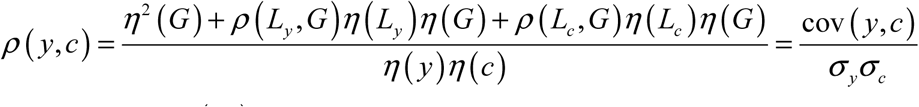

In this work, we estimate *η*^2^(*P_y_*) from the data in the following way

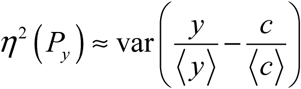

where angle brackets indicate an average over the cell population of the enclosed quantity. The validity of this estimate may not be obvious at first, so we derive it below.

Using the variance sum law, split the right-hand side of the previous equation into

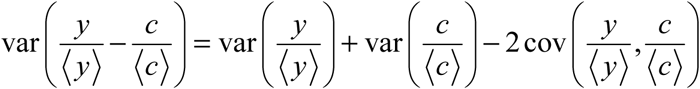

From the definition of *η*^2^, and properties of covariance, this can be re-written as

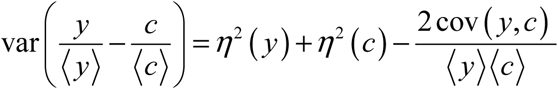

Using the definition of *η* we can say

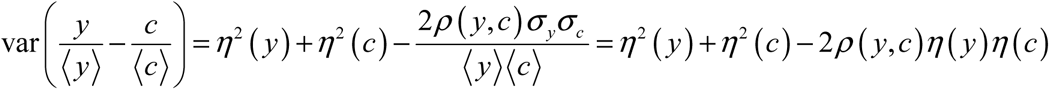

Insert the definitions from Colman-Lerner et al 2005, as above, and perform the cancellations:

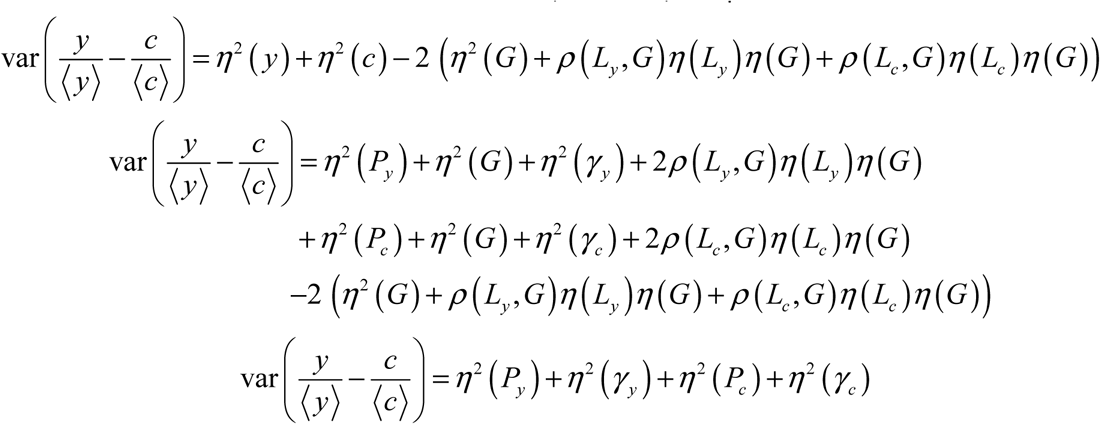

Since the constitutive pathway variation and the gene expression noise terms, i.e. the last three terms on the right-hand side, are all small (see supplement to Colman-Lerner et al 2005), this is a good way to estimate the inducible pathway variation *η*^2^(*P_y_*.)

The neglected terms are all positive, thus the computed quantity (the left-hand side) represents an upper limit for *η*^2^(*P_y_*), i.e.

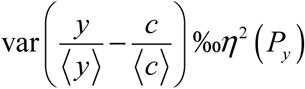

To summarize, *y* and *c* are values of the total fluorescence (two different colors) from each cell in an isogenic population. The variance of the difference between *y* and *c*, both normalized by their respective means, is a measure of the uncorrelated variation visible on a scatter plot of *y* vs. c. If both *y* and *c* are reading constitutive promoters, then this is in turn a measure of *η* ^2^ (*γ_y_*) + *η* ^2^ (*γ_c_*) + *η* ^2^ (*P_c_*), i. e. stochastic noise in gene expression. If, on the other hand, y is reading an induced promoter and *c* a constitutive promoter, then we know from previous work (Colman-Lerner et al 2005) that the variance of the difference between the normalized fluorescences includes a much larger contribution from *η*^2^(*P_y_*). We can thus use this variance as an estimate of the cell-to-cell variation in transmitted signal, *η*^2^(*P_y_*).

